# Localised negative feedback shapes genome-wide patterning of meiotic DNA breaks

**DOI:** 10.64898/2026.02.23.706156

**Authors:** Luz Maria Lopez Ruiz, Jon A. Harper, Dominic Johnson, Rachal M. Allison, William H. Gittens, George GB. Brown, Tim J. Cooper, Valerie Garcia, Matthew J. Neale

**Affiliations:** Genome Damage and Stability Centre, School of Life Sciences, University of Sussex, UK

## Abstract

Genetic diversity within sexually reproducing species arises via the formation and repair of programmed DNA double-strand breaks (DSBs) created by the evolutionarily conserved topoisomerase-like enzyme, Spo11. Because DSBs threaten genome stability, their formation is tightly regulated in both space and time. In *S. cerevisiae*, Tel1, the orthologue of mammalian Ataxia Telangiectasia Mutated (ATM) kinase, suppresses nearby DSB formation through local inhibition known as DSB interference. However, whether such local inhibition reshapes the genome-wide DSB landscape remains unclear. Here, we develop a quantitative simulation framework to model how Tel1-mediated feedback shapes Spo11-DSB formation across the yeast genome. We demonstrate that innate chromosome-specific DSB patterns, when combined with interference, generate complex, population-level redistribution of DSBs. We define the spatial range over which interference propagates and provide evidence that this regulatory mechanism requires Tel1 recruitment to DSBs via Xrs2 and Tel1 kinase activity. Although the pro-DSB factor Rec114 contributes to DSB regulation, mutation of potential Rec114 phosphorylation sites indicates that it is not an essential target of Tel1. Together, these findings demonstrate how localised negative feedback can drive broad-scale, emergent patterning of a fundamental genome-modifying process, with the potential in meiosis to influence recombination initiation and, consequently, genetic variation across generations.

## INTRODUCTION

During meiosis, a highly conserved programme of DNA double-strand break (DSB) formation and repair ensures both the accurate segregation of homologous chromosomes at the first meiotic nuclear division and the generation of recombinant allele combinations within haploid progeny^1-4^. Meiotic DSBs are catalysed by the evolutionarily conserved topoisomerase-like enzyme Spo11, the activity of which is tightly controlled by multiple, intersecting regulatory layers^5-9^. Meiotic DSBs are not randomly distributed across chromosomes. Instead, DSBs arise preferentially in relatively narrow genomic regions dispersed across the genome, termed hotspots^10^. Hotspot location is designated by a combination of species-specific factors, particularly local chromatin accessibility^8,10,11^ and, in organisms that encode PRDM9, by sequence-specific binding of this histone methyltransferase^12,13^.

In *S. cerevisiae*, DSB formation—and thus hotspot strength—is also modulated on a much broader scale via the loading of chromosome axis-associated factors, with Red1 identified as a critical component^14^. Recently, two independent modes of Red1 binding have been elucidated, mediated either by the loading and redistribution of cohesin (as marked by the meiosis-specific kleisin subunit, Rec8), or by recruitment of the HORMA-domain containing protein, Hop1, via its chromatin-binding PHD finger domain^15^. Loading of Hop1, Rec8 and Red1 enables recruitment of the pro-DSB factors Rec114, Mer2 and Mei4 (“RMM”), which have direct roles in interaction and activation of the Spo11 core complex (Spo11, Rec102, Rec104 and Ski8). The Spo11 core complex directly catalyses DSB formation. RMM components are proposed to form intermolecular condensates that may provide a catalytic surface upon which DSB formation takes place^16,17^. The localised assembly of such structures may also underscore the competition that arises at a population level between strong DSB-active regions^8,18,19^.

DSB formation is also modulated reactively. DSBs activate the evolutionarily conserved DNA damage response (DDR) kinases, Ataxia Telangiectasia Mutated (ATM) and AT-related (ATR) (Tel1 and Mec1 in *S. cerevisiae*), that regulate not only those steps critical for DSB repair such as DNA resection and inter-homologue-directed recombination^20-22^ but also help prevent premature exit into the meiotic nuclear divisions before all DSB repair has completed^7,23,24^. In *S. cerevisiae*, such transient arrest in meiotic prophase is mediated via the Mek1 kinase (the meiosis-specific Rad53 and CHK2 orthologue) and inhibition of Ndt80, the transcriptional master regulator of meiotic prophase exit^25^. Such DSB-dependent checkpoint activation thus homeostatically regulates total DSB formation by extending the temporal window of opportunity in which Spo11 can be active—a process that may enable DSBs to arise asynchronously across the genome, without premature entry into the nuclear division stage^26,27^.

ATM and ATR (and their orthologues) also act to directly limit total DSB formation and thus constrain meiotic recombination activity—a process that may arise from a combination of global and local inhibition^23,28-30^. In *S. cerevisiae*, Tel1 and Mec1 inhibit DSB formation locally, both directly in cis on the same chromatid, and in trans on the sister chromatid and homologous chromosome^16,23,31^—a collective process termed we refer to as DSB interference^7,26,31^. Such interference is proposed to arise in a DSB-dependent manner, thereby limiting the potential for DSBs to arise in close proximity within any given chromosome^7,31^. In parallel to Tel1- and Mec1-dependent pathways, DSB formation is also restricted by the homologue engagement regulatory circuit, whereby successful synapsis and inter-homologue interactions feed back to suppress further DSB formation^32^. This feedback ensures that DSB activity ceases once homologous pairing is established.

Notably, in each *S. cerevisiae* meiotic cell, around 150-200 Spo11 DSBs form^10,16,33^, equivalent to approximately one break per 250 kb of the replicated diploid genome—a density substantially lower than that of the ∼15,000 hotspots themselves (∼1 hotspot every ∼3 kb of the replicated diploid genome). Thus, in any given meiosis, only a small fraction (∼1%) of hotspots are actually active, and as a consequence each cell will generate its own unique pattern of Spo11 DSBs. Nevertheless, processes influencing DSB formation in individual cells may still shape the observed population-average frequencies of Spo11 activity—as has been previously explored^18,26^. Understanding precisely how each pathway of DSB regulation comes together to produce genome-wide patterns of DSBs remains an important challenge.

Here, we have characterised how Tel1-dependent reactive inhibition influences the chromosome-wide distribution of Spo11-DSBs. We previously established that deletion of *TEL1* leads to unique chromosome-specific spatial changes in the strength of individual hotspots^26^. We hypothesise that these patterns of change arise from Tel1-dependent DSB interference acting reactively around each DSB. To test this idea, we have developed a computational simulation of DSB interference to explore how reactive regulation can shape genome-wide DSB patterns—a process driven by the non-uniform distribution of Spo11 activity across chromosomes. We utilize strains bearing a deletion of *NDT80* and *SAE2* (the orthologue of mammalian CtIP and required to activate Mre11-nuclease-dependent release of Spo11 from DSB ends^34^; to enable total Spo11-DSB activity to be measured across the genome, and identify DSB interference parameters that best reproduce the effects exerted by Tel1 inhibition *in vivo*. We characterise the *in vivo* effects of mutants that alter the chromosome-wide DSB distribution. Finally, we employ these methods to test the involvement of key upstream mediators and downstream targets of Tel1-dependent DSB interference.

## RESULTS

### Tel1 exerts a unique spatial effect on the genome-wide distribution of Spo11-induced DSBs

To characterise genome-wide effects of DSB interference on Spo11 activity, we analysed patterns of Spo11-induced DSBs mapped in *S. cerevisiae* using CC-seq^35,36^ in the presence and absence of *TEL1*^26^. In order to reliably detect patterns of DSBs across chromosomes, analyses were performed on datasets collected from *sae2*Δ cells, in which unresected DSBs accumulate (**Fig. 1a**)^37^. To prevent confounding differences in meiotic prophase duration caused by Tel1-dependent checkpoint activation^26^, both *TEL1* and *tel1Δ* strains also carried a deletion in the *NDT80* transcription factor^38^ to arrest cells uniformly in late prophase I.

**Figure 1.**
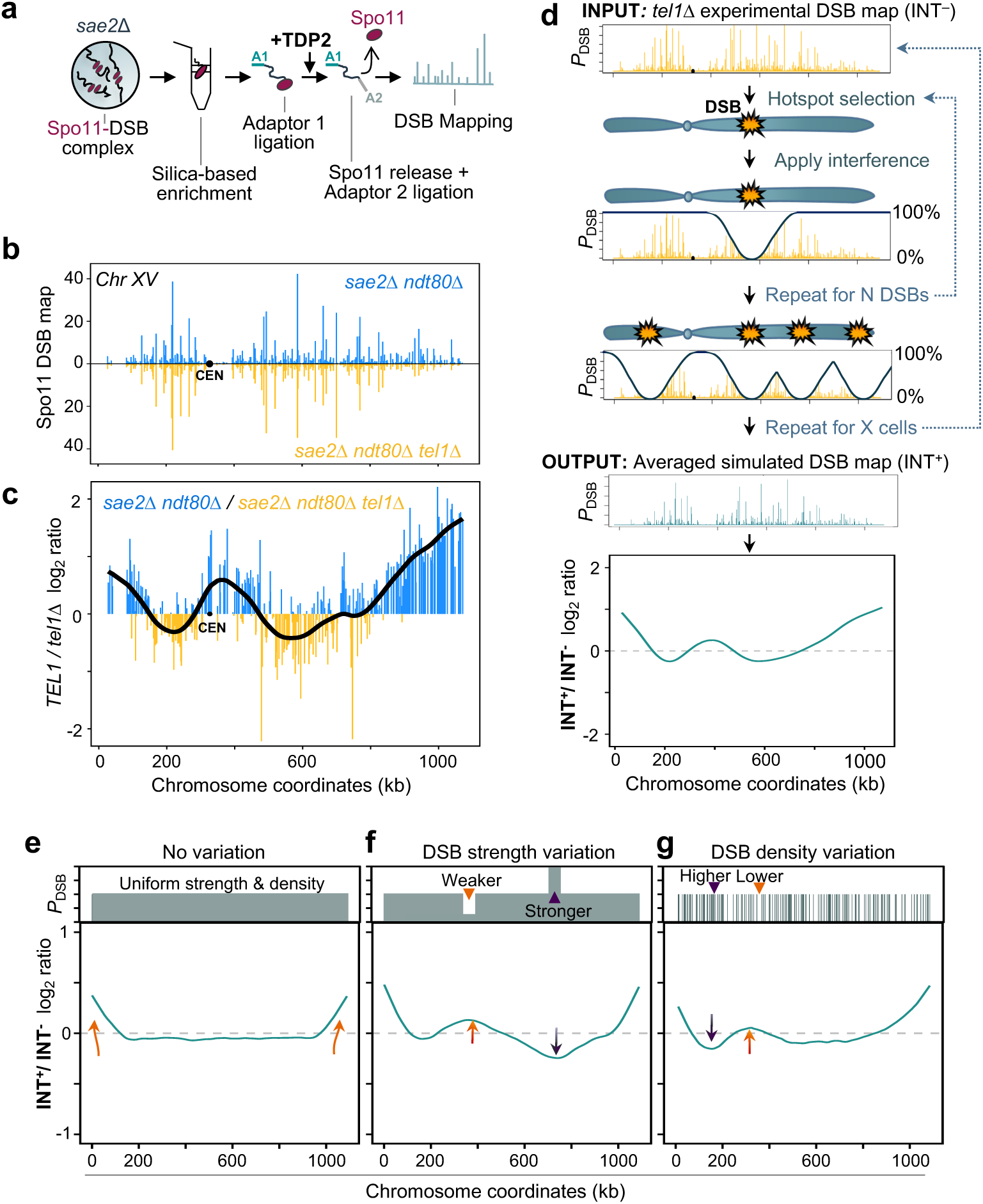
The effect of Tel1-dependent DSB interference. **a**, Schematic of the genome-wide CC-seq Spo11-DSB mapping technique (see methods). **b**, Visualization of the relative Spo11 hotspot intensities on chromosome XV in the indicated strains (NormHpChr). **c**, Log_2_ ratio of relative Spo11 hotspot intensities (NormHpChr) ± *TEL1* on chromosome XV. Black line, smoothed ± *TEL1* fold change. CEN = centromere position. Log_2_ hotspot ratio of all chromosomes are presented in **Fig. S1c**. **d**, Schematic of the interference simulator pipeline. In any given iteration of the simulation, zones of DSB inhibition are applied sequentially around each selected DSB, chosen in a weighted-random manner based on the input DSB profile (*tel1*Δ experimental input map, **INT-**). Interference is modelled following a Hann shape of decay of a specified inhibitory width. The strength of interference ranges from 100%–0% at that simulated genomic location and decays with distance. The results are aggregated (OUTPUT Averaged simulated DSB map, **INT+**) and represented by the ratio between the averaged simulated DSB map, **INT+**, and the input *tel1*Δ experimental DSB map, **INT-**. **e**–**g**, Simulation of a Hann-shaped interference window of 500 kb inhibitory width over artificial chromosome patterns of varying hotspot strengths (top panels) and the resulting +INTerference / -INTerference log2 ratio (bottom panels) when utilizing maps with (**e**) uniform position and frequency of DSBs, (**f**) regions of weak and strong hotspots inserted, (**g**) randomised distribution of hotspots of uniform strength with areas of low and high DSB density. Orange arrows indicate regions of greatest DSB formation in the presence of interference (**INT+ output**) and blue arrows indicate regions of greatest DSB formation in the absence of interference (**INT- input**) (lower panel). *P*_DSB_, probability of the DSB site. In all instances, the simulation loops through a user-specified number of DSBs (N DSBs) and iterates until a total of 10 million DSBs have been simulated.

DSB activity was quantified by summing CC-seq signal within defined hotspots across each chromosome and expressing this as a fraction of the total genome-wide signal (Normalised Hits per million reads per Chromosome, NormHpChr; see Methods) (**Fig. 1b**). As previously noted, deletion of *TEL1* caused only subtle changes in the strength of DSB formation within individual hotspots (**Fig. 1b, S1a-b**)^26^. However, when DSB distributions were expressed as the ratio of *TEL1 / tel1*Δ signal, clear spatial domains of concerted change emerged—clusters of adjacent hotspots that showed coordinated increases (blue peaks) or decreases (yellow peaks) in relative DSB activity in the presence or absence of Tel1, respectively (**Fig. 1c, Fig. S1c**). This ratio is negatively correlated with hotspot strength, suggesting that the activity of Tel1 leads to suppression of DSB formation in stronger areas, either directly or indirectly (**Fig. S1d**).

Notably, these Tel1-dependent domains were distinct and unique for each chromosome (**Fig. S1c**), suggesting that the effect of Tel1 is influenced by the underlying chromosome-specific distribution of Spo11 activity. Central chromosomal regions tended to exhibit relatively higher DSB activity in the absence of Tel1 (yellow peaks), whereas the ends of chromosomes tended to show stronger activity in the presence of Tel1 (blue peaks). This trend was less evident, or entirely absent, on the shorter chromosomes (I, VI, III, and IX) (**Fig. S1c**). Overall, these observations suggest that Tel1 modulates DSB formation in a domain-specific manner, as has been previously proposed^18,26^.

### Variation in DSB hotspot distributions alters the population-average effects of DSB interference

We hypothesised that the chromosome-specific changes caused by *TEL1* deletion arise because Tel1-dependent interference acts differently on each chromosome: depending on the number, position, and strength of their hotspots. To test this idea, we developed a simulator that models the effect of sequentially forming multiple DSBs on a chromosome in the presence and absence of interference in a large population of cells (**Fig. 1d, Fig. S2a** and **Methods**). Briefly, the simulator uses DSB maps from *sae2*Δ *ndt80*Δ *tel1*Δ (lacking interference effects) to weight the probability of a given site being selected to form a DSB. Thus, strong hotspots are more likely to be selected for DSB formation than weaker hotspots. Interference is then modelled as a decaying wave spreading from the site of a DSB, reducing the probability of further DSBs forming in sites proximal to an existing DSB (**Fig. 1d** and **Fig. S2a**). By iterating through thousands of simulated meiotic cells, a population-average DSB map is generated (INT+), in which the effects of interference can be compared to the input map (no interference, INT–) by plotting a ratio (**Fig. 1d**, lower panel).

To conceptually explore how chromosome-specific hotspot distributions can influence the effect that interference has on DSB distributions, we first ran simulations with idealised patterns of hotspot strengths, modelling interference as a Hann window of inhibitory width covering a 500 kb distance (**Fig. 1e-g** and **Fig. S2b**). When the DSB hotspot distribution was entirely uniform in strength and position (**Fig. 1e**, top), the resulting effect of interference along the chromosome was relatively flat, except for a terminal enrichment of DSBs (**Fig. 1e**, bottom). By contrast, regions of weaker or stronger hotspot strengths or densities caused localised fluctuations in the resulting DSB map, with weaker or lower density regions generally being enriched for DSBs whereas stronger or higher density regions were suppressed for DSBs (**Fig. 1f-g**).

### Simulation of DSB interference reproduces the Tel1-dependent spatial effect

Because simulated changes in hotspot strength and density produced complex interference patterns, we hypothesized that combining real hotspot maps with interference might reproduce Tel1’s effects observed in vivo. To test this, we simulated DSB formation using real hotspot data from chromosome XV as a representative example. Because the range of DSB inhibition caused by interference is unknown, we tested Hann-shaped interference windows of varying widths (10–1000 kb; **Fig. S2c**).

Applying very narrow (e.g. 50 kb) interference widths around each successful DSB generated very flat effects of interference (**Fig. S2d**, bottom), indicating minimal difference between the simulated output and the experimental *tel1*Δ DSB distribution. By contrast, intermediate interference widths (e.g. ∼300 kb) generated domains of concerted change that resembled the experimental *±TEL1* ratio pattern (**Fig. S2d**, middle), indicating that the population-average effect of Tel1 observed in vivo can be reproduced by this simple model of DSB interference. Finally, broad windows (∼900 kb) captured the enrichment at chromosome ends observed in the experimental data but less effectively reproduced finer internal structure (**Fig. S2d**, top).

In our basic model, DSB interference is only applied in cis—inhibiting DSB formation only on the same chromatid around where the DSB has formed. However, it has been suggested that DSB inhibition may arise both in cis and in trans, mediated by the complementary activities of Tel1 and Mec1^7,23,31^ (**Fig. S2e**). We tested this idea by fixing interference width at 500 kb and varying the strength of trans interference relative to fixed cis interference strength, again using chromosome XV as a representative example (**Fig. S2f**). Increasing levels of trans interference increased the amplitude but not the overall pattern of interference, indicating that trans interference may be an important consideration in our model.

Having established these parameters, we systematically tested combinations of trans interference strength and window widths (**Fig. 2a**). Because the number of DSBs in our *sae2*Δ *ndt80*Δ strain is estimated to be in the range of 150-200 DSBs^10,16,26,33^, simulations were run until 175 DSBs had formed in each virtual meiosis. Deviations between simulated and experimental *TEL1/tel1*Δ ratios were quantified by root mean squared deviation (RMSD) and plotted as a heatmap (**Fig. 2a**). A cluster of well fitting simulations (low deviation between the experimental data and simulation output) was found centred on a width of ∼500 kb (± 100 kb) and trans strength of ∼0.6–0.8 (**Fig. 2a-b**).

**Figure 2.**
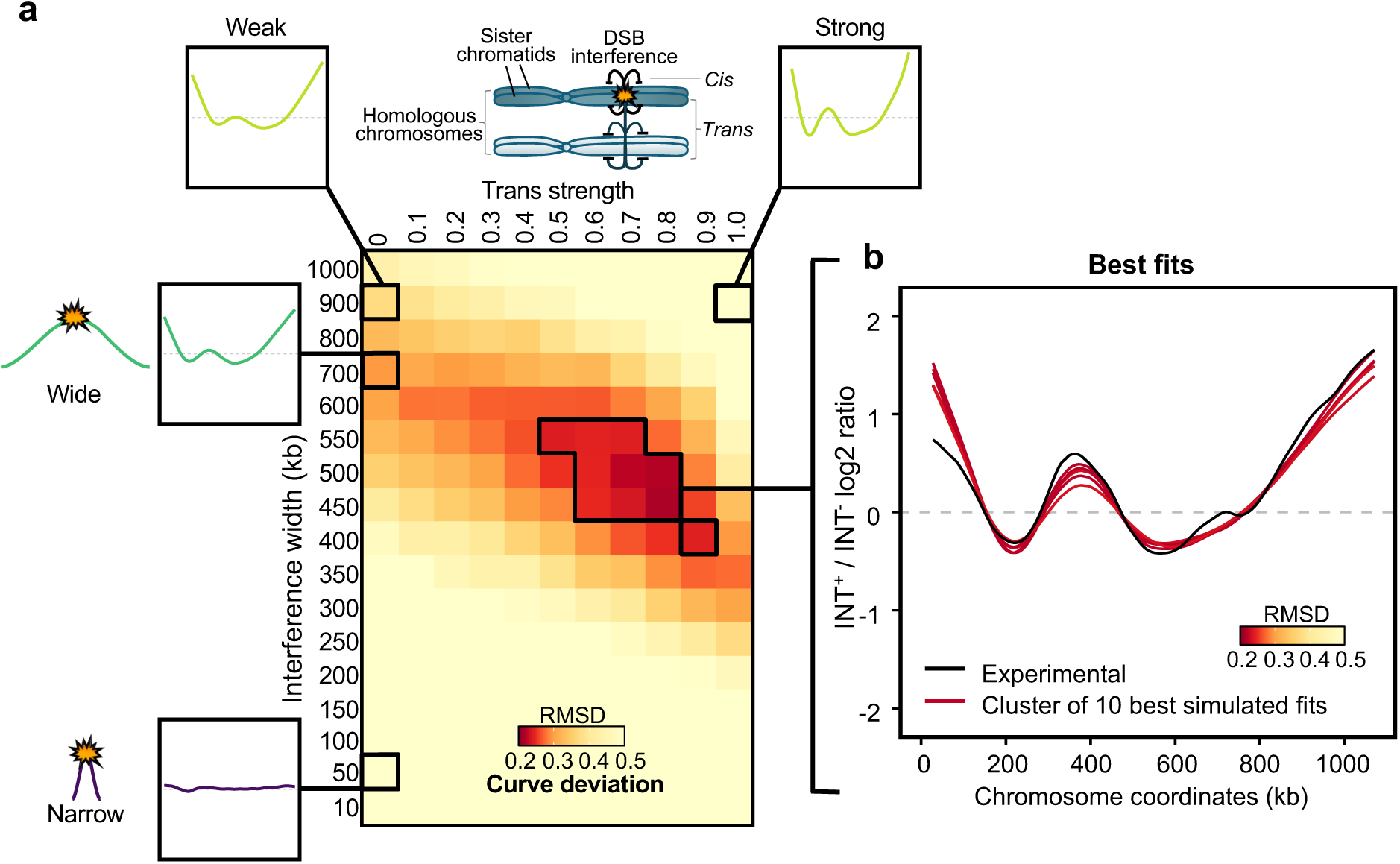
A simple model of DSB interference can accurately recapture the experimental Tel1 spatial effect. **a**, Heatmap, where each pixel represents the deviation between the smoothed fold changes observed experimentally (± Tel1 ratio), for each combination of simulated interference window width (vertical axis) and variable trans interference strength (0–1, horizontal axis) for Chromosome XV. Dark red and yellow pixels indicate areas of low (<0.2) and high (>0.5) deviation between the experimental data and the simulations, respectively. Inset plots are representative simulated INT+/INT- log_2_ ratio curves, with parameters matching indicated pixels. **b**, Comparison of the ten simulations with lowest RMSD (coloured by RMSD as in (**a**)) against the ±*TEL1* experimental smoothed pattern (black line). 175 DSBs were simulated per cell, with enough independent cells simulated to equal 10 million DSBs in total.

Our initial interference model used a Hann-shaped decay function, creating a strong inhibitory peak at the DSB site that decreased steadily with distance (**Fig. S3a ii**). Simulations using three other interference shapes (modelling different rates of decay from softer to stronger inhibition from the DSB site: Exponential, Tukey 0.5, and Tukey 0) were also capable of producing results that resembled experimental data, but with slightly different parameter combinations (**Fig. S3a-c**). Notably, windows where the interference remained locally strong but then rapidly decayed over shorter distances (e.g. Tukey 0.5, and Tukey 0; **Fig. S3a iii-iv**) produced best-fitting simulations with narrower window widths (∼200–300 kb) and weaker trans interference strengths (0.2–0.6) than Hann windows (**Fig. S3b-c ii-iv**). By contrast, exponentially decaying windows (**Fig. S3a i**) required narrow interference windows (∼280 µ) and the greatest strength of trans interference (0.8–1.0) to produce the best fits (**Fig. S3b-c i**).

Overall, although all decay shapes were able to generate good fits, the effect of interference produced from simulations using the Hann window yielded the lowest deviation from the experimental *TEL1/tel1*Δ ratio (**Fig. S3b-c ii**). Thus, to streamline further analysis, we henceforth used Hann windows as a model of interference effects. On chromosome XV, simulations utilising a Hann window of 400-550 kb wide and trans strength of 0.6-0.9 were our best-fitting model of the Tel1-dependent effects observed *in vivo*. Simulating lower or higher numbers of DSBs also produced good fits, and with near-identical window widths, but with modest compensatory shifts in the strength of trans interference (**Fig. S4**). Importantly, because of the decaying shape and symmetry of the interference functions, we emphasise that substantial (>50 %) interference only reaches a fraction of the stated distance (**Fig. S2b**). i.e. Even with a Hann window of 500 kb, a DSB arising just 125 kb from another will only be subject to a 50% reduction in probability (**Fig. S2b**). A >80% reduction in probability only arises when a second DSB is within ∼75 kb of another (**Fig. S2b**). Such distances are thus broadly consistent with locus-specific measures in which the effects of DSB interference were not detected beyond ∼150 kb^31^.

### Effects of Tel1 on DSB distributions across different chromosomes can be recaptured by simulation

To assess whether the principles of Tel1-dependent interference identified on chromosome XV also apply genome-wide, we extended our simulations to all sixteen *S. cerevisiae* chromosomes using the Hann window shape of decay (**Fig. 3a**). Overall, the simulation reproduced the chromosome-specific Tel1-dependent DSB patterns observed experimentally on large and medium chromosomes (>500 kb) generating clustered areas of low deviation between simulated and experimental ±*TEL1* fold-change profiles, indicating robust fits across a range of parameter values (**Fig. 3a**, upper panels). Visual inspection of the ten best-fit simulations per chromosome confirmed that these models captured the qualitative, domain-level patterning unique to each chromosome (**Fig. 3a**, lower panels).

**Figure 3.**
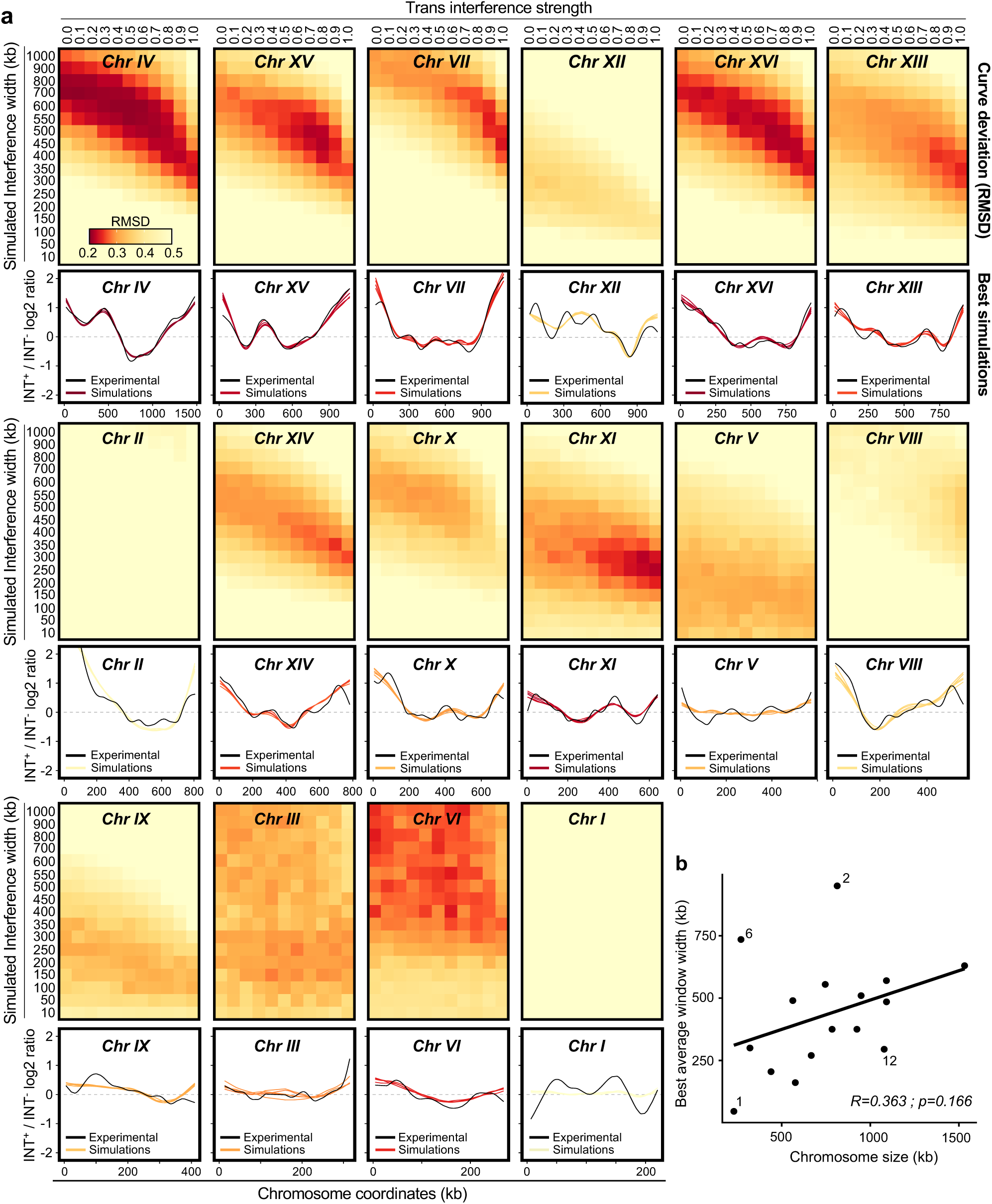
Simulation of DSB interference accurately recaptures the effect of Tel1 across *S. cerevisiae* chromosomes. **a**, (Upper panels) Heatmaps, where each pixel represents the deviation between the smoothed fold changes observed experimentally (±*TEL1* ratio) for each combination of simulated interference window width (vertical axis) and variable trans interference strength (0–1, horizontal axis) across the sixteen *S. cerevisiae* chromosomes. Dark red and yellow pixels indicate areas of low (<0.2) and high (>0.5) deviation between the experimental and the simulations, respectively. (Lower panels) Comparison of the simulations with lowest RMSD (coloured by RMSD as in the upper panels) against the ±*TEL1* experimental smoothed pattern (black line). 175 DSBs were simulated per cell, with enough independent cells simulated to equal 10 million DSBs in total. **b**, Correlation between the window width of best fitting simulations (averaged window width of the 10 simulations with the lowest RMSD) against chromosome size.

In contrast, simulating interference on the three shortest chromosomes (<400 kb) did not clearly identify clustered areas of low deviation (**Fig. 3a**, bottom). In particular, chromosomes III and VI showed scattered and weak fits across multiple window widths. We note that the shorter chromosomes generally lack major regions of fold change ±*TEL1* (**Fig. S1c**), which may limit the ability of the simulation to generate meaningful visual and statistical matches (**Fig. 3a**, bottom). Simulations of chromosomes II and XII were fitted less well by simulation despite being relatively large, potentially due to complications arising from the chromosomal deletion of *TEL1* on chromosome II and the rDNA which is located in the middle of chromosome XII. Interestingly, we noticed a positive trend between interference width and chromosome length (R=0.363 and *p*=0.166; **Fig. 3b**) that was slightly increased upon removing these two chromosomes (R=0.398 and *p*=0.08), suggesting that the spread of Tel1-dependent DSB inhibition may differ by chromosome length.

Collectively, these simulations demonstrate that a simple model of local reactive DSB inhibition can reproduce the complex chromosome-specific patterns of DSB activity characteristic of Tel1-dependent DSB interference.

### Altering the biological distribution of Spo11 DSBs alters the signature of interference

To further verify that the signature of Tel1-dependent DSB interference is shaped by the underlying genomic distribution of Spo11 activity, we genetically modified the hotspot landscape using a Gal4BD–Spo11 fusion protein (*SPO11-GBD*), which additionally directs Spo11 to otherwise cold genomic regions containing Gal4 upstream activating sequences (UAS) (**Fig. 4a**)^19,39^. As expected, CC-seq maps confirmed the formation of novel DSB hotspots at UAS sites, including in regions normally cold for Spo11 activity (**Fig. 4b**; red arrow), substantially modifying the regular distribution of Spo11 activity when assessed at a chromosome-wide scale (R=0.61; **Fig. S5a**).

**Figure 4.**
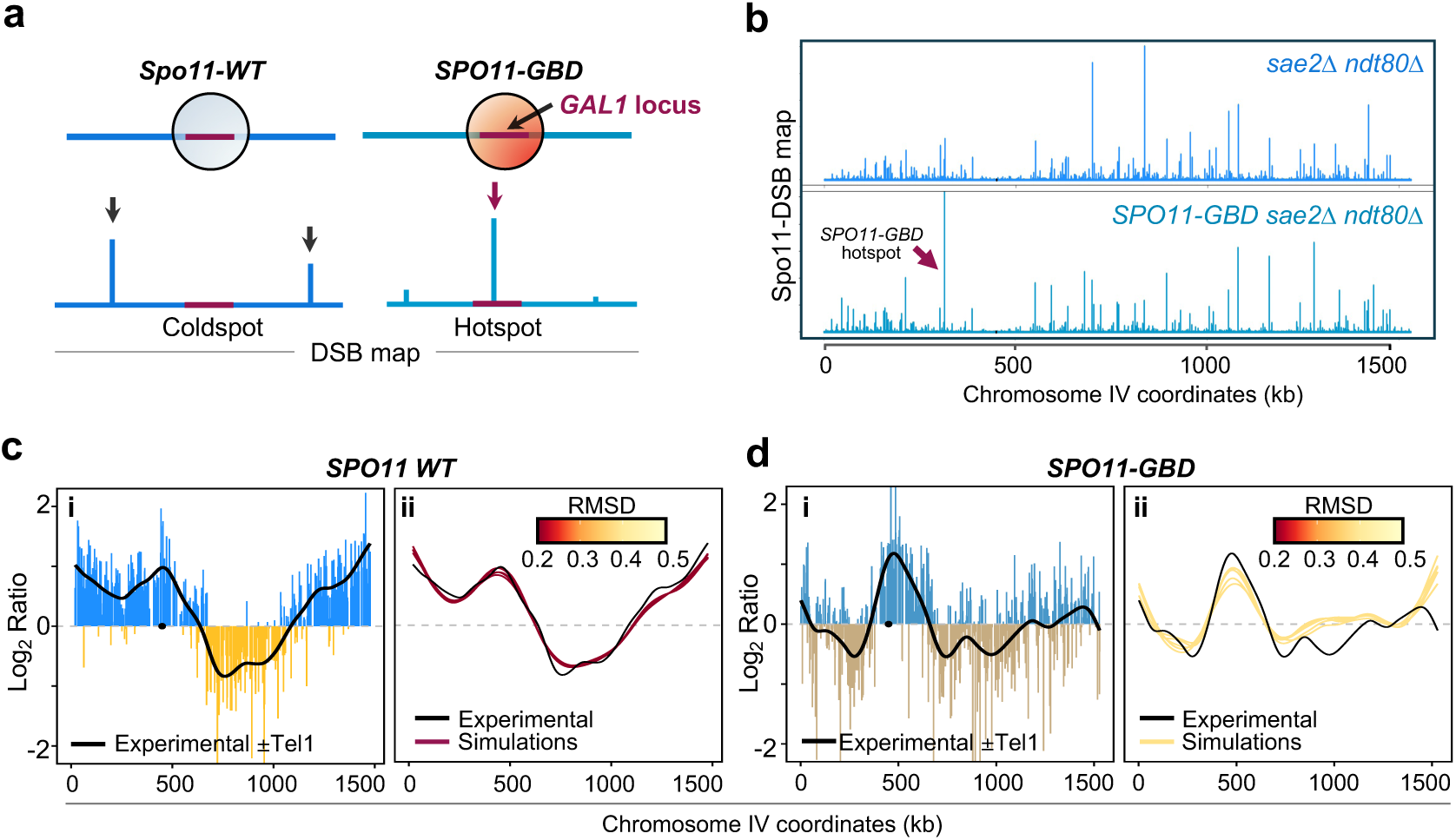
Altering the biological distribution of Spo11 DSBs generates new signatures of interference. **a**, Representative cartoon of the Spo11-GBD system. A Gal4BD-Spo11 fusion protein (*Spo11-GBD*) is expressed to additionally target DSB formation in regions of the genome that contain a Gal4 binding motif (UAS) (as developed by Robine et al., 2007). **b**, Visualization of the relative Spo11 hotspot intensities (NormHpChr) on chromosome IV in the indicated strains. The purple arrow indicates the position and frequency of a strong Spo11-GBD specific hotspot. Other chromosomes are presented in **Fig. S5d. c-d**, **i** Ratio of relative Spo11 hotspot intensities ±*TEL1* on chromosome IV in the SPO11-WT (**c**) and SPO11-GBD (**d**) systems. Values above zero indicate a higher DSB frequency in the presence of Tel1 and below zero a higher DSB frequency in the absence of Tel1. Fold change was smoothed to highlight the spatial trend effect of *TEL1* deletion (black line) in these backgrounds. Other chromosomes are presented in **Fig. S5e**. **ii** Representation of the simulations (INT+/INT- log_2_ ratio) that best recaptures the experimental *±TEL1* smoothed pattern (NormHpChr log2 ratio) in the SPO11-WT (**c**) and SPO11-GBD (**d**) backgrounds in chromosome IV. Lines are coloured by deviation (RMSD) between the experimental and the simulated Log_2_ ratios. Other chromosomes are presented in **Fig. S5f**.

Similar to the *SPO11* wild-type background, within the *SPO11-GBD* system, deletion of *TEL1* caused a relatively modest global change (R=0.94; **Fig. S5a**). However, the spatial pattern of these Tel1-dependent changes was distinct: regions of enhanced and reduced hotspot activity occurred at novel positions along the chromosomes, reflecting the altered hotspot map in the *SPO11-GBD* strain (**Fig. 4c–d i** and **Fig S5b–e**). To assess whether these changes could again be explained by local DSB interference, simulations were performed on the five largest chromosomes using the *SPO11-GBD tel1*Δ dataset as input. Such simulations were able to reproduce the altered experimental *±TEL1* fold-change displayed within the *SPO11-GBD* system (**Fig. 4c–d ii** and **Fig. S5f**).

These findings demonstrate that modifying the biological pattern of Spo11-DSB activity alters the Tel1-dependent interference signature, reinforcing our findings across different *S. cerevisiae* chromosomes, and again suggesting that Tel1 acts through a local, reactive mechanism with population-average effects that are governed by the unique strength and spatial distribution of DSB hotspots present on each chromosome.

### Tel1 kinase activity is required for Tel1-dependent DSB interference

Previous studies indicate that Tel1 kinase activity contributes to the global suppression of meiotic DSB formation^18^. However, whether the effect of Tel1 on DSB interference is kinase-dependent has not been directly tested. Given that Tel1-dependent interference underlies the population-average fold changes in DSB hotspot strengths we reasoned that, if such regulation depends on Tel1’s kinase activity, a kinase-dead mutant (*tel1-kd*) should display fold-change patterns comparable to those observed in *tel1*Δ cells.

To test this idea, we mapped Spo11 DSBs genome-wide in a *sae2*Δ *ndt80*Δ *tel1-kd* strain using CC-seq. Inactivation of the Tel1 kinase activity revealed a spatial pattern of hotspot redistribution very similar to that observed in *tel1*Δ cells (R=0.97) (**Figure 5a-b** and **Fig. S6a-c**), supporting the view that the effect that Tel1 causes is dependent on its kinase activity. Moreover, as expected given the similarity of the *tel1*Δ and *tel1-kd* phenotypes, interference simulations were able to model the interference lost in the *tel1-kd* dataset—now also including good fits for chromosome II, the site of the *TEL1* gene, which is not deleted in the *tel1-kd* strain (**Fig. 5c** and **Fig. S7a**). A positive (even stronger) correlation between chromosome length and best-fitting interference width was again observed (R=0.682, p=0.0036; **Fig. S7b**). Together, these results indicate that Tel1 kinase activity is required for DSB interference, again supporting a model in which Tel1’s catalytic activity locally inhibits further DSB formation near existing breaks.

**Figure 5.**
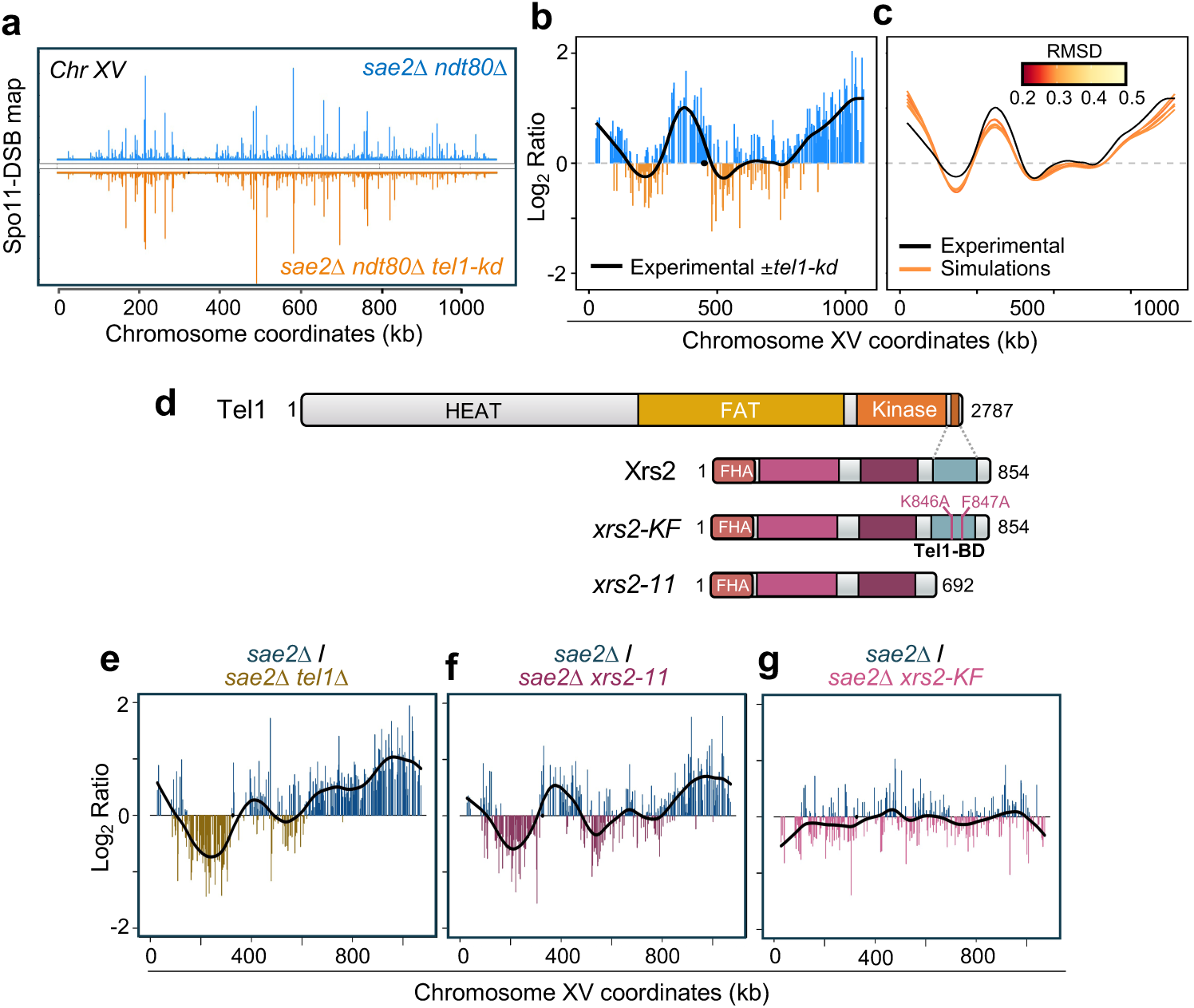
Tel1 kinase activity and the Xrs2 C-terminal domain are required to mediate DSB interference. **a**, Visualization of the relative Spo11 hotspot intensities (NormHpChr) on chromosome XV in the indicated strains. **b**, Ratio of relative Spo11 hotspot intensities (NormHpChr log2 ratio) ±*tel1-kd* on chromosome XV. Values above zero (blue peaks) indicate a higher DSB frequency in the presence of Tel1 and below zero (orange peaks) a higher DSB frequency in the *tel1-kd* mutant, respectively. Fold change was smoothed to highlight the spatial trend effect of the *tel1-kd* mutant (black line). All chromosomes are presented in **Fig. S6b**. **c**, Representation of the simulations (INT+/INT- log_2_ ratio) that best recapture the experimental ±*tel1-kd* smoothed pattern (NormHpChr log_2_ ratio) in chromosome XV. Lines are coloured by deviation (RMSD) between the experimental and the simulated Log2 ratios. Other chromosomes are presented in **Fig. S7a**. **d**, Domain structure of Tel1, Xrs2 and Xrs2 mutants. Key residues in the Tel1 binding domain of Xrs2 are highlighted. **e-g**, Ratio of relative Spo11 hotspot intensities (NormHpChr log_2_ ratio) ±*TEL1* (**e**), ±*xrs2-11*(**f**) and ±*xrs2-KF* (**g**) on chromosome XV. Values above zero indicate a higher DSB frequency in the presence of Tel1 (**e**) or Xrs2 WT (**f-g**) and below zero a higher DSB frequency in the absence of Tel1 (**e**), *xrs2-11* (**f**) or ±*xrs2-KF* (**g**). Fold change was smoothed to highlight the spatial trend effect of *TEL1* or *XRS2* as indicated (black line). Other chromosomes are presented in **Fig. S8c**.

### The C-terminal domain of Xrs2 is required for Tel1-dependent DSB interference

Recent studies have demonstrated that, in meiotic cells, Xrs2 plays a critical role in recruiting Tel1 to chromosomal loop–axis contact sites, where DSBs are proposed to be initiated^40^. Specifically, the C-terminal domain of Xrs2 appears to mediate Tel1 binding, positioning Tel1 to locally inhibit further DSB formation in nearby chromosomal regions. To directly test whether this Tel1 recruitment mechanism is also required for the genome-wide interference effects observed here, we examined two C-terminal Xrs2 mutants—*xrs2-11* containing a truncation of 162 amino acids at its C-terminus^41^, and *xrs2-KF* containing two amino acid changes (K846A/F847;^42^)—that disrupt Tel1 interaction with DSBs to different degrees (**Fig. 5d**)^40^.

First, to test whether either strain displayed known characteristics similar to *TEL1* deletion, we examined a known Tel1-suppressed hotspot within the *YCR061W* gene^26^. Interestingly, this hotspot was markedly elevated in *xrs2-11*, but not in the *xrs2-KF* mutant (**Fig. S8a**) supporting the idea that there may be differential Tel1 impairment between the two mutants. Consistent with this idea, global inter-hotspot correlations (**Fig. S8b**) and chromosome-wide fold-change analyses revealed that *xrs2-11* displayed spatial patterns of DSB redistribution similar to those observed in *tel1*Δ *cells* (though slightly attenuated in amplitude), whereas *xrs2-KF* showed no clear pattern relative to the control (**Fig. 5e-g** and **Fig. S8c**). Taken together, these results indicate that the C-terminal domain of Xrs2 is likely critical for Tel1-dependent DSB interference, but that the *xrs2-KF* point mutations are insufficient to abolish this function. Thus, we infer that, even though checkpoint deficient^40^, *xrs2-KF* retains sufficient Tel1 recruitment and/or activation to sustain substantial interference between Spo11-induced DSBs in meiosis.

### Rec114 is not an essential target of DSB interference

Our results indicate that Tel1 kinase activity is essential for DSB interference, yet the critical downstream targets of this regulation remain unclear. A prime candidate is Rec114—an evolutionarily conserved factor required for Spo11-DSB formation^30,43^. Prior work in both *S. cerevisiae* and *C. elegans* support the idea that phosphorylation of Rec114 (Dsb-1 in *C. elegans*) by Tel1 or ATM, respectively, inhibits its pro-DSB activity, thereby reducing further DSB formation^30,44,45^.

To explore whether Rec114 phosphorylation is essential to mediate DSB interference, we mapped DSB patterns in two previously established Rec114 mutants^30^: *rec114-8A*, in which eight putative phosphorylation sites are mutated to alanine (preventing phosphorylation and thus potentially phenocopying a constitutively active state), and *rec114-8D,* in which these residues are replaced with aspartic acid (mimicking phosphorylation and thus a constitutively inactive state; **Fig. 6a**). If Tel1-mediated phosphorylation of Rec114 is required for interference, we would predict that *rec114-8A*, the non-phosphorylatable mutant, would mimic *tel1*Δ while *rec114-8D* would not. Contrary to this expectation, hotspot strengths were similar in the *sae2*Δ *rec114-8A* mutant as the *sae2*Δ control (*R=*0.95; **Fig. S9a**), and no clear chromosomal fold-change pattern was observed (**Fig. 6b** and **Fig. S9b**). By contrast, the *sae2*Δ *rec114-8D* mutant displayed a substantially altered DSB landscape (*R=*0.83; **Fig. S9a**), and chromosomal fold-change patterns in *rec114-8D* partially resembled those in *tel1*Δ (**Fig. 6b** and **Fig. S9b**), albeit with almost as much difference to the *sae2*Δ *tel1*Δ strain (R = 0.86) as to the *sae2*Δ control (R = 0.83) (**Fig. S9a**).

**Figure 6.**
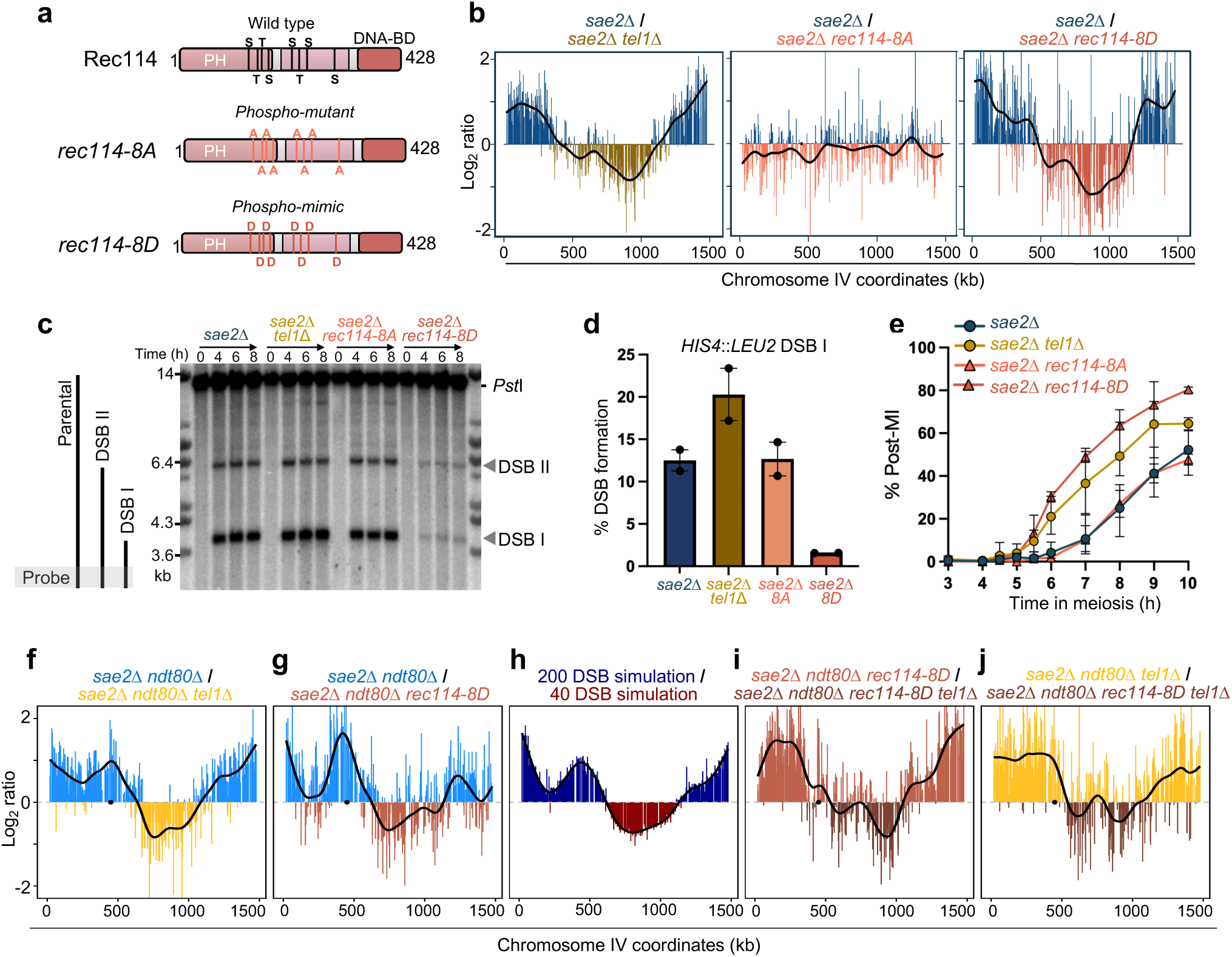
Rec114 is not an essential target of DSB interference. **a**, Domain structure of Rec114. Key putative phosphorylation target residues are highlighted. **b,** Ratio of relative Spo11 hotspot intensities (NormHpChr log_2_ ratio) ±*TEL1* (left panel), ±*rec114-8A* (middle panel) or ±*rec114-8D* (right panel) on chromosome IV. Values above zero indicate a higher DSB frequency in the presence of Tel1 (left panel) or Rec114 wild type (middle and right panels) and below zero a higher DSB frequency in the absence of Tel1 (left panel) or in the presence of *rec114-8A* (middle panel) or *rec114-8D* (right panel) constructs. Fold change was smoothed to highlight the spatial trend effect of *TEL1* deletion or the *rec114-8A/8D* mutants (black line). Other chromosomes are presented in **Fig. S9b**. **c**, Representative Southern blot of *Pst*I digested genomic DNA isolated at the specified times, hybridised with *MXR2* probe at the *HIS4*::*LEU2* locus. The lower grey filled triangle indicates the quantified DSB (DSB I). **d**, Quantification of DSB I (average of 6–8 h time points, n=2). Error bars indicate range (averaged 6–8 hours time points). Points represent individual repeats. **e**, Meiotic nuclear division (MI and MII) kinetics were assessed by counting the appearance of bi-, tri- and tetra-nucleate DAPI-stained cells. At least 100–200 cells were scored for each timepoint after inducing meiosis entry. Averaged biological replicates of n=4 for *sae2*Δ and *sae2*Δ *tel1*Δ, and n=2 for *sae2*Δ *rec114-8A and sae2*Δ *rec114-8D* are represented. Error bars represent range. **f-g**, Ratio of relative Spo11 hotspot intensities (NormHpChr log_2_ ratio) ±*TEL1* (**f**) and ±*rec114-8D* (**g**) in the *ndt80*Δ background on chromosome IV. Values above zero indicate a higher DSB frequency in the presence of Tel1 (**f**) and Rec114 wild type (**g**) and below zero a higher DSB frequency in the absence of Tel1 (**f**) or presence of *rec114-8D* (**g**). Fold change was smoothed to highlight the spatial trend effect of *tel1*Δ *or rec114-8D* as indicated (black line). **h**, Ratio of relative Spo11 hotspot intensities from DSB interference simulations performed with DSB frequencies of 200 and 40. Interference applied with a 200 kb width following a Hann shape of decay in cis on chromosome IV. In each instance, iterative independent simulations were repeated until a total of 10 million DSBs had been simulated in each condition. **i-j**, Ratio of relative Spo11 hotspot intensities (NormHpChr log_2_ ratio) ±*TEL1* (**i**) and ±*rec114-8D* (**j**) in the indicated background on chromosome IV. Values above zero indicate a higher DSB frequency in the presence of Tel1 (**i**) or *rec114-8D* (**j**). Fold change was smoothed to highlight the spatial trend effect of *tel1*Δ *or rec114-8D* as indicated (black line). Other chromosomes are presented in **Fig. S9d**.

Taken together, these results suggest that the phosphomimetic, and not the phospho-mutant, allele displays features somewhat consistent with abrogated DSB interference. More importantly, the lack of any effect in the *rec114-8A* mutant indicates that Rec114 is not an essential target of Tel1 for mediating the interference effect.

### Rec114 modulates DSB distributions independently of Tel1

Unlike the *rec114-8A* allele, the *rec114-8D* allele has defects in Spo11-DSB formation^30^. Given the surprising results observed in the *REC114* mutants, we hypothesised that at least part of the *rec114-8D*-dependent changes in DSB distributions may be caused by the lower DSB frequency. In keeping with previously reported findings, Southern blotting at the *HIS4::LEU2* locus confirmed that DSB formation in the *rec114-8D*—but not the *rec114-8A*—strain was substantially reduced (**Fig. 6c-d**)^30^. Additionally, the meiotic prophase checkpoint present in *sae2*Δ cells was bypassed even more effectively in the *rec114-8D* mutant than by *TEL1* deletion, consistent with low DSB frequency (**Fig. 6e**).

Because premature exit from meiotic prophase—caused by inefficient Tel1-dependent checkpoint activation—can itself alter spatial patterns of Spo11-DSB formation^26^, we re-examined the effect of the *rec114-8D* mutant in the *ndt80*Δ background. Notably, the *sae2*Δ *ndt80*Δ *rec114-8D* strain, whilst continuing to bear some similarity to the effect of deleting *TEL1* (R=0.91) remained equally different to the control (R=0.90; **Fig. 6f-g** and **Fig. S9c**), indicating that differences in DSB distribution caused by *rec114-8D* are neither caused by differential prophase kinetics nor identical to loss of Tel1 activity.

We reasoned that reduced DSB frequency may partially mimic the effects of reduced interference because, if fewer DSBs are created, less interference will result. To test this hypothesis, the DSB simulator was used to model the effect of varying overall DSB frequency in the presence of interference. From a ratio of high (200) and low (40) simulated DSB frequencies, a fold-change pattern was observed that was similar to the effect of *TEL1* deletion, with the emergence of domains of relative enrichment and depletion (**Fig. 6h**), thus supporting our hypothesis. Nevertheless, if the effects of *rec114-8D* were entirely due to reduced interference caused by a reduction in DSB frequency, then we would expect to observe little effect of *TEL1* deletion in a *rec114-8D* background. To directly test this idea, we compared DSB distributions between *sae2*Δ *ndt80*Δ *rec114-8D* and *sae2*Δ *ndt80*Δ *rec114-8D tel1*Δ cells (**Fig. 6i** and **Fig. S9d**). Counter to this expectation, clear spatial differences remained, indicating that Tel1-dependent effects remain active in *rec114-8D* cells despite the reduction in DSB frequency. Moreover, the amplitude of the *TEL1-*dependent effect was similar in magnitude to the effect observed in *REC114* wild-type cells, suggesting that the effect of Tel1 is independent of not just Rec114, but potentially also independent of DSB formation itself (**Fig. 6i**).

Collectively, our observations support the view that at least some of the effects of *rec114-8D* are independent of DSB interference—further supported by the observation that changes in DSB distributions caused by *rec114-8D* are independent of *TEL1* deletion (**Fig. 6j**). Thus we conclude that although changes in the effect of DSB interference may arise indirectly from globally reduced Spo11 activity, such an effect is not sufficient to explain differences in DSB distribution observed in *rec114-8D* mutants.

## DISCUSSION

Previous studies in meiotic *S. cerevisiae* cells revealed that Tel1 enforces distance-dependent interference between closely spaced Spo11-DSBs, a mechanism of DSB regulation potentially conserved in other eukaryotes^26,28,29,31,44,46^. Nonetheless, it remained uncertain over what distances the effects of interference could manifest, and whether such local control within individual cells could influence Spo11-DSB distributions observed across cell populations. Available maps of Spo11-oligo intermediates in wild-type cells appeared to support only a limited contribution towards such regulation^18^.

A major obstacle to determining the extent to which interference can influence DSB formation was the reliance on low-throughput assays that measured interference at a small number of selected hotspots^31^. We began work to overcome this limitation by developing a genome-wide assay that captures Spo11-DSBs that accumulate when Mre11 nuclease activity is inhibited, revealing strong Tel1-dependent changes in DSB distribution that we hypothesised were caused by DSB interference^26^. Here, we have developed and employed a model of Tel1-dependent interference in single cells to test this idea, estimate distances over which interference may act, and to explain the population-level effects that DSB interference may cause (**Fig. 7**). Our findings build upon previous evidence of a role for Tel1 in DSB interference^26,31^. By applying a probabilistic model of interference to a *tel1*Δ DSB map across individual simulated meioses, population-level patterns of Tel1 action can be recaptured on a genome-wide scale, suggesting that previously described effects of Tel1 on DSB interference characterised at single loci are indeed occurring across the entire genome^31^.

**Figure 7.**
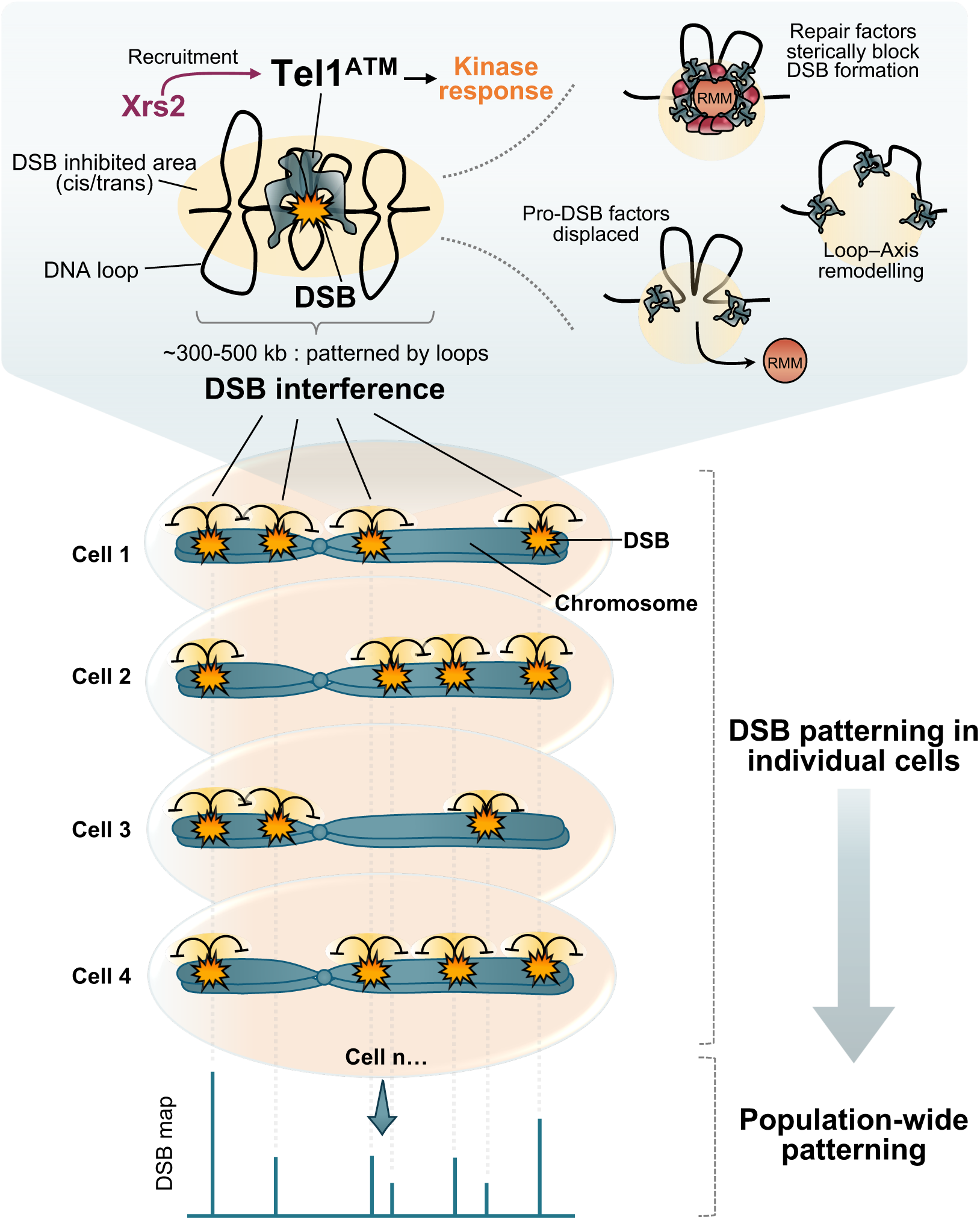
Local inhibition gives rise to distributional changes at the population level. Tel1-dependent DSB interference is reliant upon the C terminal of Xrs2 and kinase activity of Tel1. Local inhibition of DSBs results in breaks being distributed evenly across individual chromosomes at the cell level. Such distributional changes manifest in population averaged maps of Spo11 DSB activity. We propose that Tel1 kinase activity mediates these effects in individual cells by phosphorylating one or more targets in a largely DSB-dependent manner. We speculate that such reactive localised negative feedback on DSB formation could arise via recruitment of repair factors around break sites, eviction of pro-DSB factors (e.g. Rec114, Mer2, Mei4; RMM), and/or remodelling of loop–axis interactions—any or all of which will reduce subsequent Spo11 catalytic activity in the vicinity of an existing DSB.

An intrinsic assumption of how our simulation framework functions is that DSB locations are selected based on the weighted strengths of each hotspot—a relationship which means that, on average, DSBs in stronger hotspots are more likely to be selected earlier. Whilst this relationship seems quite rational, it does not have to be the case. Thus, the fact that simulations are able to recreate the effects of interference observed in vivo suggests that this intrinsic relationship is likely to be generally true. Nevertheless, we consider it likely that this relationship is not universal, and thus may underpin some of the remaining deviations that we observe between experimental and simulation datasets.

It is notable that the effects of Tel1 manifest differently on each chromosome—with clear domainal changes visible at different locations. Whilst such changes might have been explained by some upstream change in DSB potential prior to break formation, our simulations support the view that such domainal changes arise reactively due to the inherent differences in hotspot positions and strengths that exist on each chromosome. One simple feature of interference is the relative enrichment of DSB formation towards chromosome ends. We attribute this to the fact that such distal chromosome regions are less likely to be subject to an interfering signal than are hotspots located within the central areas. Weaker regions also receive relatively more DSBs in the presence of interference, suggesting that one evolutionary benefit of this process may be to more evenly spread Spo11 activity across the genome.

Nevertheless, whilst simulations were able to recapture these chromosome-specific effects, they could not do so without some variation in parameters—most notably the width over which the decaying interference signal was modelled, which displayed a positive trend with chromosome size in both our *tel1*Δ and *tel1-kd* simulations (**Fig. 3b** and **Fig. S7b**). One possibility to explain this effect could be differential (greater) compaction of chromatin on larger chromosomes in three-dimensional space causing relative increases in the spread and propagation of the interfering signal. Thus, such differences in compaction could explain why larger chromosomes were better fit with simulations that utilized wider windows of interference, while, in general, narrower windows better suited smaller chromosomes. If these effects prove robust and broadly applicable, the precise range of interference is therefore likely to differ not just between chromosomes, but also perhaps between species based on their average DNA loop size and the degree of chromosome compaction during prophase^47^. Loops may also underpin the apparent best-fitting shape of the decay rate of interference. The Hann window, with its relatively strong degree of inhibition immediately around the DSB may mimic the ‘averaged’ effect of chromosome compaction, where variably sized loops and/or adjacent loop clusters are the unit upon which Tel1-dependent inhibition acts^8^ (**Fig. 7**).

Application of interference in trans—between chromatids—was also critical for improving model fits. Whilst it was not possible to determine a universal strength of trans interference for all chromosomes, simulations nonetheless support the view that there is at least some degree of suppression of Spo11 activity on a second chromatid that is not subject to breakage, as has been proposed^23^. Due to the lack of downstream steps of recombination in our experimental set up, we favour the view that the trans interference we are modelling is between sister chromatids^8,31^. Consequently, due to the potential lack of trans interference between homologous chromosomes, estimates of trans interference strength may be lower than in the natural system. However, such a difference is likely to be offset by the persistence of DSBs (and therefore interference) in *sae2*Δ cells (see below).

It is important to consider the mechanism by which the apparent negative regulation of Spo11 activity is being applied. We have provided supporting evidence that DSB interference is dependent on recruitment of Tel1 to DSB sites via the C-terminal domain of Xrs2^40^, and is dependent on Tel1 kinase activity. In principle, Spo11 and/or any number of the factors it interacts with^48,49^ could be direct targets of phospho-regulation. Tel1-dependent recruitment of repair factors could either physically block Spo11 from binding nearby, or could lead to the exclusion of pro-DSB-factor binding close to an active DSB (**Fig. 7**). Alternatively, given the important role chromatin accessibility plays on Spo11-DSB formation^10^, it could be the local chromatin structure, or indeed its higher-order conformation—for example loop–axis interactions (**Fig. 7**)^8,50^—that are subject to regulation.

Our experimental work seems to rule out phosphorylation of Rec114 as an essential component. Nevertheless, the complex consequences of the hyper-negatively charged *rec114-8D* allele point to a compound modulation of the normal regulatory pathway whereby Tel1-dependent effects seem to arise even under conditions of reduced DSB activity. In *C. elegans*, the Rec114 orthologue Dsb-1 behaves as if it is indeed a bona fide target of the ATM kinase^44,45^. Whether this indicates deviation from a shared regulatory pathway involving Rec114, or differing genetic redundancies across species remains to be determined. More importantly, the critical target(s) for Tel1-dependent DSB interference in *S. cerevisiae* at present remains unsolved.

Interestingly, hotspot strength was negatively correlated with both the ±*TEL1* ratio (**Fig. S1d**) and the ±*rec114-8D* ratio (**Fig. S9e**), suggesting that weaker hotspots are promoted by both Tel1 and Rec114. This similarity in effect, in addition to the reduced DSB frequency observed in *rec114-8D* strains, may explain why *rec114-8D* appears to partially mimic a loss of DSB interference, but for very different reasons: DSB interference indirectly promotes DSB formation in weaker hotspot regions, whereas *rec114-8D* is predicted to have a particularly pronounced and direct defect in forming DSBs at weaker hotspots because it is a DSB-formation hypomorph.

It is important to note that the effects of Tel1 deletion on DSB distributions described here were not detectable in the *SAE2+* background^18,26^. Whilst we cannot exclude that the methodological differences used in mapping DSBs (Spo11-oligo-seq vs CC-seq) are the source of this difference, we favour the view that the apparent lack of similar population-average effects of *TEL1* deletion in the presence of Sae2 are because DSB interference is short-lived and ceases upon DSB repair. Thus, as a result of this shorter life-span, interference could be limited in both longevity and spread in *SAE2+* cells, whilst nonetheless still ensuring that DSBs in two adjacent hotspots rarely exist at the same time^31^.

Processes that arise downstream of DSB repair—such as synapsis-dependent DSB shut off^32,51^—may also act to overwrite and obscure the influence DSB interference has on DSB distributions. It may also be that the persistence of unresected DSBs in the *sae2*Δ background leads to heightened effects of Tel1-dependent interference. Notably, pilot studies in *rad50S* and *mre11-H125N* strains—which also both accumulate unprocessed DSBs^52,53^—lead to effects similar to *sae2*Δ, suggesting that if this is the case, it is not a consequence of Tel1 hyperactivity that is unique to *sae2*Δ cells^24^. More likely, we favour the view that in wild-type cells Tel1 plays a substantial role locally in suppressing coincident DSB formation^18,31^, handing over to other stages of regulation once resection and repair begins. With this in mind, it seems possible that Mec1 may complement and/or substitute for Tel1 at later stages, regulating DSB formation both within and between chromatids^23^. Such a model would predict larger population-average effects to manifest whenever there are defects and/or delays in efficient repair—something that could happen stochastically from cell to cell, and also variably between genomic loci *within* a cell. Thus regardless of whether DSB interference is constrained spatially or temporally, this does not preclude the possibility of DSB interference (whether mediated by Tel1 or other factors) playing an important role in the positioning and regulation of DSB formation.

### Outlook

As the vital first step for meiotic recombination, DSB formation and regulation remain important subjects for study. Our observations provide insight into how processes at the site of a single DSB can shape genome-wide patterns of DSB formation, and therefore how they might dictate the positions of subsequent recombination events observed in a population. Notably, one effect of this regulation is to indirectly encourage DSB formation in weaker hotspot areas, and suppress DSBs in hotter areas—an effect felt across all chromosomes, and that may have potential ramifications for downstream processes such as crossover formation. Looking more broadly, our results demonstrate how localised, individual effects are capable of building broad population-level impacts. Specifically, whatever the biological process, for anything that is patterned in space and time, the negative feedback associated with localised reactive event-dependent inhibition will indirectly promote activity in regions that would otherwise be unlikely to receive such events. Thus, our observations are not limited to studies of DSB patterning, instead being potentially applicable to a wide range of biological systems and concepts that are subject to localised reactive regulation.

## METHODS

### Yeast strains

All *Saccharomyces cerevisiae* yeast strains used in this study (**Supplementary Table 1**) are in the SK1 background and derived using standard techniques. Strains contain the *his4X*::*LEU2* and *leu2*::*hisG* exogenous sequences inserted on chromosome III^16,35^, and carry the *ndt80*Δ*::LEU2, tel1*Δ::*HphMX4* and/or *sae2*Δ::*kanMX* gene disruption alleles^20,27,31,38^. The *rec114-8A* and *rec114-8D* alleles are as described^30^, with amino-acid substitutions re-validated by sequencing. The Gal4BD-Spo11 fusion allele is as described^39^. All integrations were confirmed by DNA amplification from genomic DNA or colony PCR. The *xrs2-11* and *xrs2-KF* (K846A/F847A) alleles are as described^40-42^.

### Culture methods

For meiosis induction, a single colony was inoculated in 4 mL of YPD medium (1% yeast extract, 2% peptone, 2% glucose supplemented with 0.5 mM adenine and 0.4 mM uracil) and incubated at 30 °C, 250 rpm for ∼24 hours to reach saturation, then diluted to OD600 of 0.2 either YPA (1% yeast extract, 2% peptone, 1% potassium acetate) or SPS (0.5% yeast extract, 1% peptone, 0.67% Yeast Nitrogen Base without amino acids, 1% potassium acetate, 0.05M Potassium Hydrogen phthalate, 0.001% Antifoam 204) pre-sporulation medium. Cultures were incubated at 30 °C, 250 rpm for 14–16 hours, then washed and resuspended in pre-warmed SPM sporulation medium (2% potassium acetate supplemented with diluted amino acids) and incubated at 30 °C, 250 rpm for the duration of the time course. Samples were taken at the relevant timepoints and processed differently. For DNA extraction, 1 mL samples were taken at t = 0, 4, 6 and 8 hours after inducing meiosis. Samples were centrifuged at 3000 x *g* for 4 minutes, and pellet resuspended in 50 mM EDTA, centrifuged again for 1 minute at 3000 x *g*, and pellet stored at -20 °C until use. For Spo11 CC-seq libraries, 50 mL of samples were taken at t = 6 hours, spin at 3000 x *g* for 5 minutes and pellets frozen at -20 °C until use. For FACS, samples were taken at t = 0, 2, 4, 6 and 8 hours after inducing meiosis, samples were centrifuged at 16,000 x *g* for 1 minute, fixed in 70% EtOH and stored at 4 °C until use. For DAPI staining, samples were taken at t = 3, 4, 5, 5.5, 6, 7, 8, 9 and 10 hours after inducing meiosis. Cells were fixed in 100% EtOH and stored at -20 °C until use.

### FACS

To monitor entry into meiotic prophase, samples were centrifuged at room temperature, 16,000 x *g* for 1 minute, pellet resuspended in 10 mM Tris HCl pH 8.0 / 15 mM NaCl / 10 mM EDTA pH 8.0 / 1 mg/Ml RNase A and incubated at 37 °C for 2 hours at 800 rpm. Samples were then centrifuged at 16,000 x *g* for 1 minute, pellets resuspended in 1 mg/mL Proteinase K + 50 mM Tris HCl pH 8.0 and incubated at 50 °C for 30 minutes at 800 rpm on. Samples were spun, pellets were washed in 1M Tris-HCl pH 8.0 and then resuspended in 50 mM Tris-HCl pH 8.0 + 1 µM Sytox green. Samples were stored overnight at 4 °C and then sonicated at 20% amplitude for 12–14 seconds before being sorted by flow cytometry (Accuri™ Flow Cytometers).

### Cell fixation and DAPI staining

Ethanol-fixed cells were dried at RT on a glass slide, stained with 2 µL of Fluoroshield™ DAPI Sigma-Aldrich (F6057-20ML) and 100–200 mono-, bi-, tri-, tetra-nucleate cells were scored by microscopy (Zeiss AXIO) using fluorescence (CoolLED pE-300 lite). Meiotic progression was determined based on the frequency of cells that entered MI (binucleated) or MII (tri-, tetra-nucleate) at different timepoints after inducing meiosis (t = 3, 4, 5, 5.5, 6, 7, 8, 9 and 10 hours).

### Proteolytic genomic DNA extraction

Meiotic cell culture pellets were defrosted at room temperature, resuspended in spheroplasting mix: spheroplasting buffer (1 M sorbitol / 100 mM NaHPO4 pH 7.2 / 100 mM EDTA), zymolyase 100T (50 mg/mL) and 1% β-mercaptoethanol, and incubated at 37 °C for 1 hour. Cells were lysed with 3% SDS / 0.1 M EDTA / 50 mg/mL Proteinase K, and incubated overnight at 60 °C. After cooling at room temperature for 5 minutes, proteins were removed with two rounds of phenol/chloroform separated by a 5-minute rest and 5 minutes centrifugation at 14,000 rpm. DNA and RNA were extracted from the aqueous phase and precipitated with 3 M NaAc pH 5.2 and 100% EtOH, centrifuged at 14,000 rpm for 1 minute, aspirated and washed with 70% EtOH, pulsed down, air dried for 10 minutes and resuspended in 1x TE (10 mM Tris / 1 mM EDTA pH 7.5) overnight at 4 °C. RNA was digested with 1 mg/mL RNase A for 1 hour at 37 °C. DNA was precipitated with NaAc pH 5.2 and 100% EtOH, mixed by inversion and spun for 1 min at 14,000 rpm. DNA was washed with 70% EtOH, pulsed down, air dried for 10 minutes, dissolved in 1x TE (10 mM Tris / 1 mM EDTA pH 7.5) overnight at 4 °C. To measure the frequency of DSBs genomic DNA was digested with *Pst*I restriction enzyme and incubated overnight at 37 °C.

### DSB analysis by Southern blot and hybridisation

Digested genomic DNA was separated on 0.7% 1x TAE agarose gels at 45–50 V for 15–19 hours in the presence of 0.1 mg/mL EtBr. DNA was nicked by exposure to 1800 J/m2 UV in a Stratalinker, then soaked in denaturing solution (0.5 M NaOH, 1.5 M NaCl), on a shaker for 30 minutes. Denatured gels were transferred to a Biorad Zeta-probe membrane under vacuum (50–55 mBar for ∼2 hours) in 0.5 M NaOH, 1.5 M NaCl. The membrane was washed twice with 2x SSC (300 mM NaCl, 30 mM trisodium citrate pH 7.0) and then thrice with distilled water. The membrane was cross-linked by exposure to 1200 J/m2 UV in a Stratalinker, dried at room temperature for 1 hour and stored at 4 °C until probed. Southern blot membranes were incubated with pre-warmed hybridisation solution (0.5 M NaHPO_4_ pH 7.5, 5% SDS, 1 mM EDTA, 1% BSA) at 65 °C for ∼1–2 hours. To quantify the DSBs at the *HIS4*::*LEU2* locus, membranes were hybridised with the *MXR2* DNA probe (**Supplementary Table 2**)^31^—radiolabelled with P_32_ prepared via random priming using the High Prime kit (BioRad). Pre-hybridisation solution was discarded, and the membrane incubated with hybridisation solution containing the radioactive probe overnight at 65 °C. After incubation, the membrane was washed (1% SDS / 40 mM NaHPO_4_ / 1 mM EDTA), air dried, and exposed to a phosphor screen overnight. After exposure (usually 8–48 hours), the phosphoscreen was scanned with a Fuji FLA 5000 reader and analysed using ImageGauge software (Fuji).

### Covalent complex sequencing (CC-seq) mapping

Protein-DNA Covalent-Complex mapping (CC-seq) in yeast followed a method previously described^26,35,36^. Briefly, meiotic cell samples are chilled on ice then frozen at -20 °C for at least 8 hours, then thawed and spheroplasted (1 M sorbitol, 50 mM NaHPO4, 10 mM EDTA, 30 min at 37°C), fixed in 70% ice-cold ethanol, collected by centrifugation, dried briefly, then lysed in STE (2% SDS, 0.5 M Tris, 10 mM EDTA). Genomic DNA was extracted via Phenol/Chloroform/IAA extraction (25:24:1 ratio) at room temperature, with aqueous material carefully collected, precipitated with ethanol, washed, dried, then resuspended in 1x TE buffer (10 mM Tris/1 mM EDTA). Total genomic DNA was sonicated to <500 bp average length using a Covaris M220 before equilibrating to a final concentration of 0.3 M NaCl, 0.1% TritonX100, 0.05% Sarkosyl. Covalent complexes were enriched on silica columns (Qiagen) via centrifugation, washed with TEN solution (10 mM Tris / 1 mM EDTA / 0.3 M NaCl), before eluted with TES buffer (10 mM Tris / 1 mM EDTA / 0.5% SDS). Samples were treated with Proteinase K at 50 °C for 1 hour and purified by ethanol precipitation. DNA ends were filled and repaired using NEB Ultra II end-repair module (NEB #E7645), with adapters ligated sequentially to the sonicated, then blocked, ends with recombinant TDP2 treatment in between these steps to remove the 5-phosphotyrosyl-linked Spo11 peptide. Ampure bead cleanups were used to facilitate sequential reactions. PCR-amplified libraries were quantified on a Bioanalyser and appropriately diluted and multiplexed for deep sequencing (Illumina MiSeq 2x75 bp).

FASTQ reads were aligned to the reference genome (SacCer3H4L2; which includes the *HIS4*::*LEU2* and *leu2*::*hisG* loci inserted into the Cer3 *S*. *cerevisiae* genome build^16,35,36^ via Bowtie2, using TermMapper as previously described^16,35,36^ (https://github.com/Neale-Lab/terminalMapper), with all subsequent analyses performed in R version 4.1.2 using RStudio (Version 2021.09.0 Build 351). Reproducibility between libraries for independent biological replicates was evaluated and validated prior to averaging. Details of individual libraries are presented in **Supplementary Table 3**.

### Calibration of CC-seq libraries

For each library the proportion of non-specific reads (background reads) were estimated by measuring the hit rate per million reads per base pair (HpM) in 47 of the longest gene ORFs (> 5.5 kb long) in the *S. cerevisiae* genome. This value (usually <0.01 HpM per base pair) was subtracted proportional to the width of each hotspot. Remaining hotspot signals were expressed as a proportion of total hotspot signal (NormHpM) and additionally normalised per chromosome (NormHpChr) prior to use in simulations. In *tel1-kd* cells, hotspot widths were extended by 300 bp on each side (NormHpChr300) to capture the hotspot signals that spread in this mutant.

### Hotspot identification

Hotspots were identified in our baseline *sae2*Δ *ndt80*Δ strains as described^26^. For the *SPO11-GBD* analysis a different hotspot template was created by identifying hotspots in *sae2*Δ *ndt80*Δ, *sae2*Δ *ndt80*Δ *tel1*Δ, *sae2*Δ *ndt80*Δ *SPO11-GBD and sae2*Δ *ndt80*Δ *SPO11-GBD tel1*Δ libraries.

### Simulation of Tel1 DSB interference

Simulated DSB distributions were produced using a simulator built in R (version 4.3). The simulator generates virtual chromosomes of user-specified length and number, binning coordinates by user-specified bin widths (set as 100 bp wide for all data presented here). Each bin is assigned a relative probability based on the input DSB hotspot map. For the purposes of our analysis, we used virtual chromosomes based on the *S. cerevisiae* genome.

When a DSB is generated, a window of interference is applied, decreasing the probabilities of DSB formation in adjacent bins in a manner where such repression decays with distance. Such inhibition thereby reduces or eliminates the chance that subsequent DSBs will occur in such bins.

To fit experimental data, each simulation continued for a specific chromosome until the total number of genome-wide DSBs was equivalent to that estimated to occur in our control strain (e.g. 150-200 DSBs per cell). To minimize sampling issues and emulate the population-level data of CCseq and ensure that the results presented are representative and equally powered simulations were repeated until 10 million DSBs had been simulated for each combination of conditions (e.g. interference window width, trans interference strength). Within the simulation, key parameters that can be adjusted are: 1) The decay shape (Hann, Tukey, Exponential) and width (in kb or slope; µ) of the interference window; 2) The total genome-wide DSB frequency; 3) Whether DSB interference is applied only in *cis*, or is also applied in *trans* on sister chromatids and/or homologues.

### Bioinformatic analysis of Spo11-DSBs

All bioinformatics analyses were performed in R (R version 4.1.2) using RStudio (Version 2021.09.0 Build 351). Scripts are available on https://github.com/Neale-Lab/Tel1_DSB_Interference_LLR. Raw (FASTQ) libraries are available via the GEO repository GSE245327.

### Statistical analysis

Model fits were assessed quantitatively by comparing the deviation (root mean squared deviation; RMSD) between the simulated and experimental fold-change maps. All statistical analyses were performed in R version 4.4.2.

## DATA ACCESSIBILITY

Raw FASTQ libraries are available from the GEO repository under accession numbers GSE245327 and GSE319316. Processed hotspot average table files and analysis scripts are available at https://github.com/Neale-Lab/Tel1_DSB_Interference_LLR. Processed singlet and averaged Fullmap files are available at https://figshare.com/s/41992f2861d66d26d739; 10.25377/sussex.31314022 (reserved DOI).

## Acknowledgements

M.J.N., L.M.L.R, J.A.H, R.M.A., D. J., T.J.C., G.G.B.B. and W.H.G. were supported by the Wellcome Trust Investigator Award 200843/Z/16/Z, Wellcome Trust Discovery Award 225852/Z/22/Z, and ERC consolidator grant 311336. W.H.G. was also independently supported by the Biotechnology and Biological Sciences Research Council Discovery Fellowship BB/V005081/1. We thank Antony Oliver (University of Sussex) for the gift of recombinant TDP2 catalytic domain used in our CC-seq library preparation.

## Author contributions

Conceptualisation: M.J.N., L.M.L.R., J.A.H

Data Curation: L.M.L.R, J.A.H

Formal Analysis: L.M.L.R, J.A.H., M.J.N

Funding acquisition: M.J.N., W.H.G.

Investigation: L.M.L.R., J.A.H., R.M.A., D.J., G.G.B.B., T.J.C., M.D., V.G., M.J.N.

Methodology: G.G.B.B., W.H.G., M.J.N., J.A.H., T.J.C.

Project administration: M.J.N.

Software: J.A.H., T.J.C. L.M.L.R, G.G.B.B., W.H.G., M.J.N.

Supervision: M.J.N., V.G.

Visualisation: L.M.L.R, J.A.H, M.J.N.

Writing—original draft: M.J.N., L.M.L.R, J.A.H

Writing—review and editing: M.J.N., L.M.L.R, J.A.H

---

**Supplementary Figure 1.**
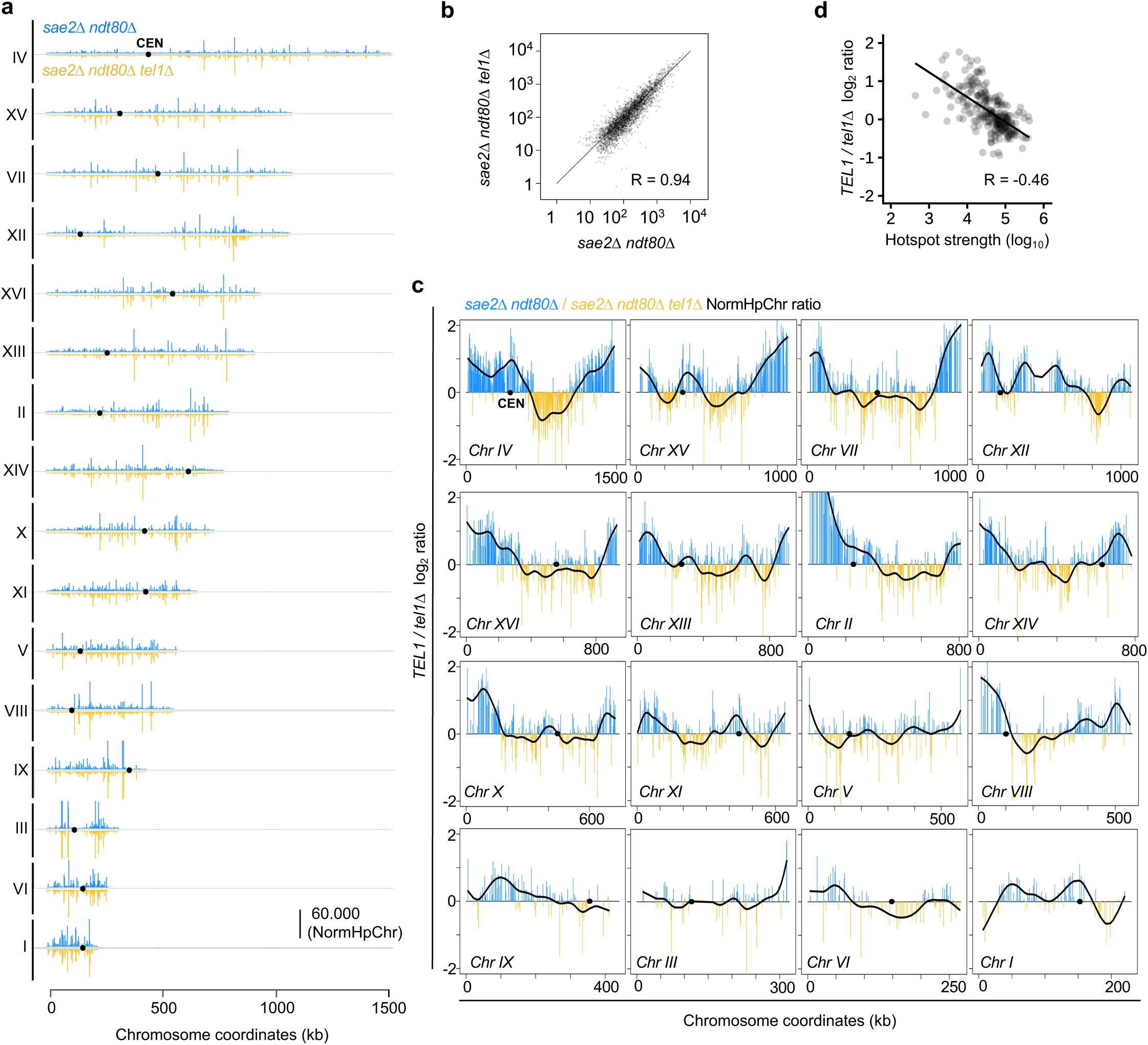
Effects of *TEL1* deletion across *S. cerevisiae* chromosomes. **a**, Visualization of the relative Spo11 hotspot intensities in the indicated strains (NormHpChr) and chromosomes. **b,** Pearson correlation in hotspot strength between the indicated strains. **c**, Log_2_ ratio of relative Spo11 hotspot intensities (NormHpChr) ±*TEL1* on each chromosome. Black line, smoothed ±*TEL1* fold change. CEN, centromere position. **d**, Pearson correlation between the *TEL1*/*tel1*Δ log_2_ ratio and Spo11 hotspot intensities across all chromosomes.

**Supplementary Figure 2.**
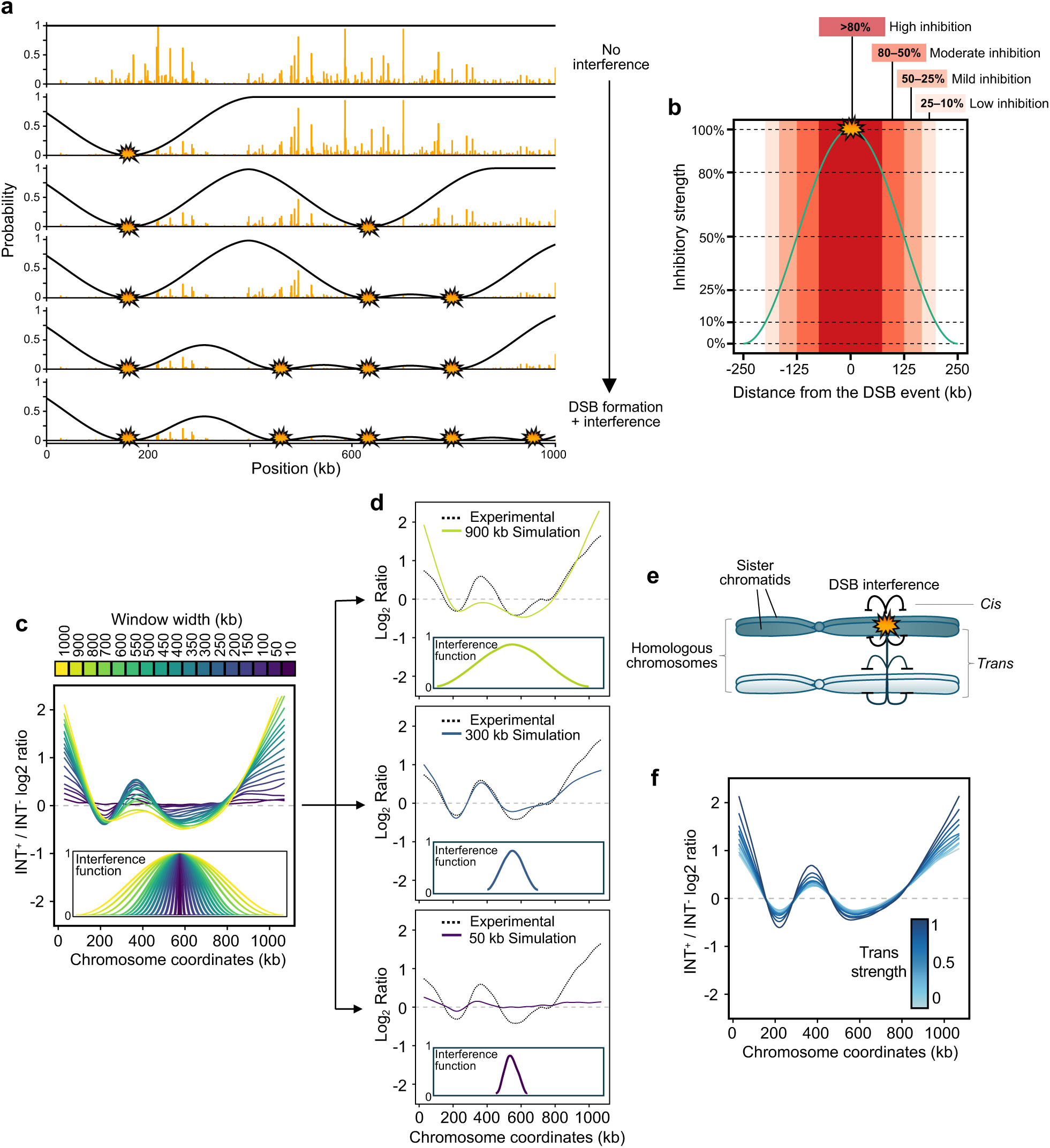
Simulation of DSB interference. **a**, Example virtual chromosome plots from representative simulation showing sequential generation of interference (500 kb Hann shape) emanating from each round of DSB formation. Each wave of interference reduces the relative probability of future DSBs from forming in flanking bins. The virtual chromosome here is 1000 kb long, equivalent to *S. cerevisiae* chromosome XV, and bins are 100 bp wide. **b**, Representation of simulated inhibition strength against distance from the DSB for a given window of interference (Hann, 500 kb wide). The inhibition strength is the proportion by which the probability of a DSB forming at a given position is reduced, with 100% inhibition reducing the probability to 0. **c**, Log_2_ ratio of simulated hotspot intensities ±interference on chromosome XV. Lines are coloured by the width of the Hann interference functions (shown inset). **d,** Selected simulations as in **c**, compared to experimental data (grey dotted line). **e**, Schematic representation of trans interference, in which DSBs inhibit the formation of DSBs on another chromatid at the same locus. The interference window is fixed at Hann 500 kb. **f**, Log_2_ ratio of simulated hotspot intensities ±interference on chromosome XV, interference window width 500 kb. Lines are coloured by the strength of trans interference relative to cis interference.

**Supplementary Figure 3.**
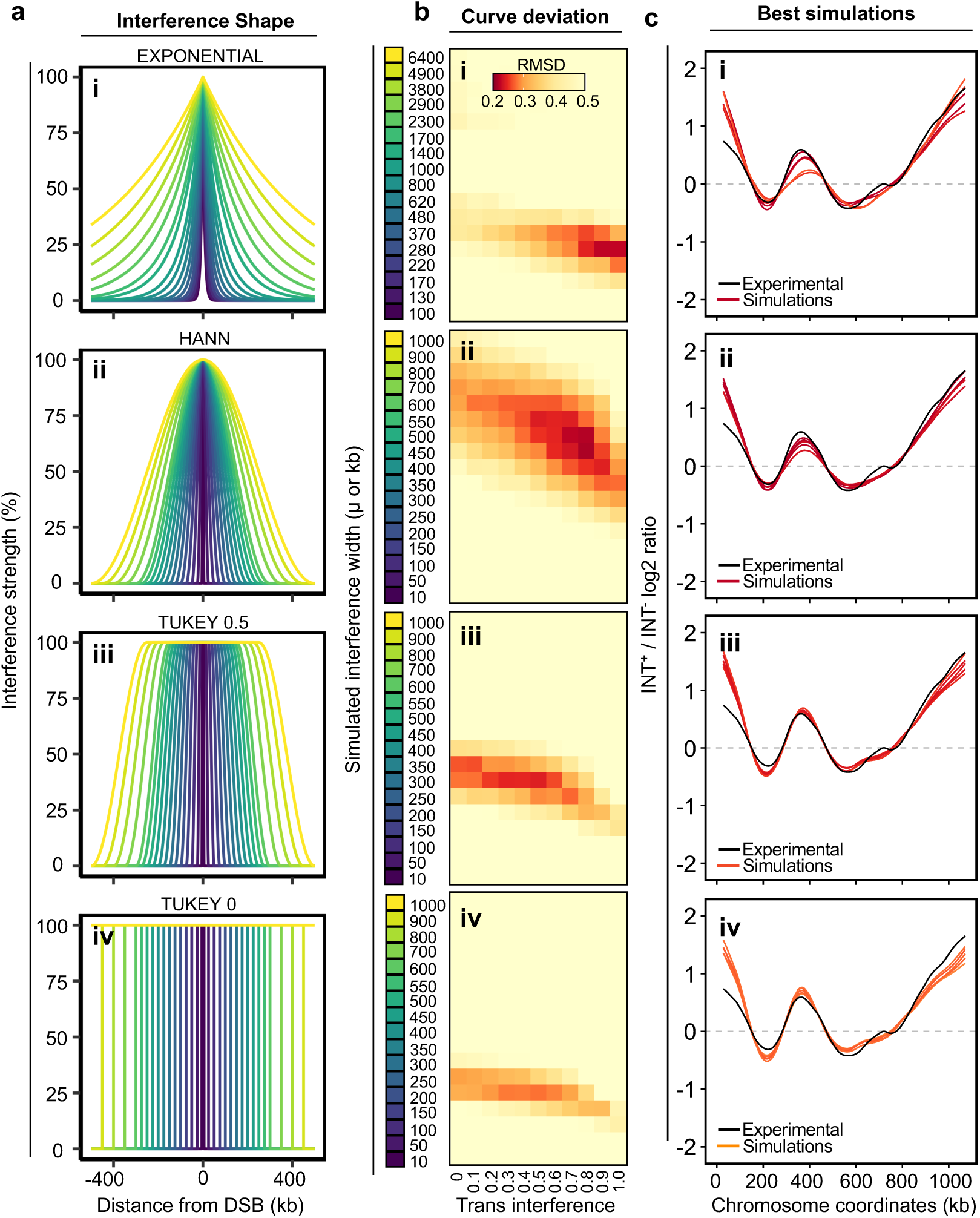
Simulations with different windows of decay. **a**, Representation of the strength and decay of inhibition (100–0%) that emanates from a simulated DSB with a given shape of interference window: Exponential (**i**), Hann (**ii**), Tukey 0.5 (**iii**) and Tukey 0 (**iv**). Plots indicate the area inhibited by varying window widths of interference ranging from 100–6400 µ for Exponential decay and 10–1000 kb for Hann, Tukey 0.5 and Tukey 0 decay. **b**, Heatmaps, where each pixel represents the deviation between the smoothed fold changes observed experimentally (±*TEL1* ratio) for each combination of simulated interference window width (vertical axis) and variable trans interference strength (0–1, horizontal axis) across the sixteen *S. cerevisiae* chromosomes. Dark red and yellow pixels indicate areas of low (<0.2) and high (>0.5) deviation between the experimental and the simulations, respectively. **c**, Comparison of the simulations with lowest RMSD (coloured by RMSD as in (**b**)) against the ±*TEL1* experimental smoothed pattern (black line). 175 DSBs were simulated per cell.

**Supplementary Figure 4.**
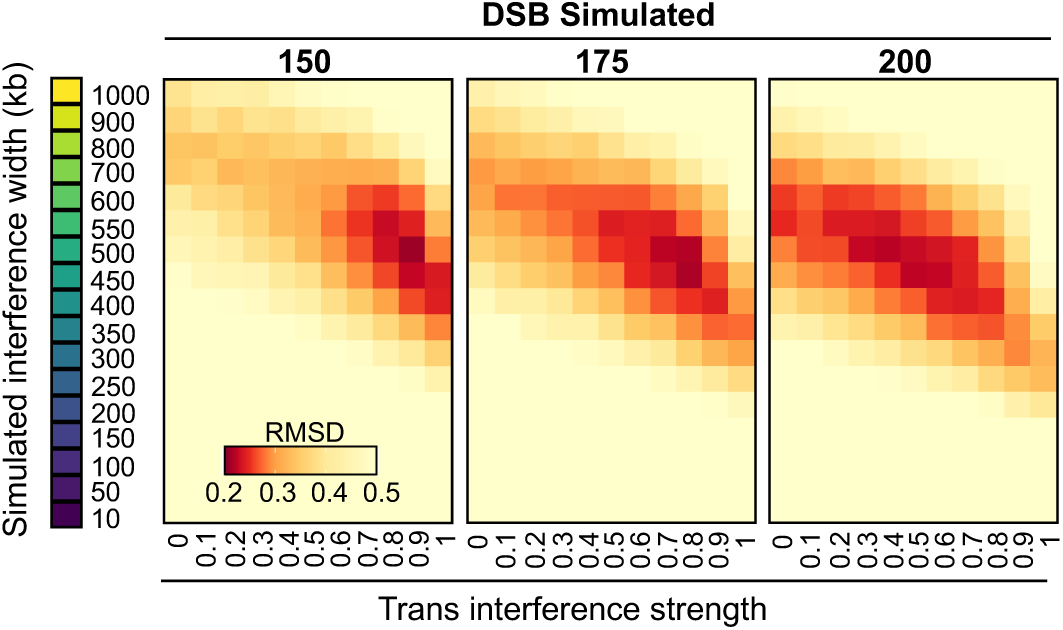
Effect of simulated DSB number on fits to experimental data. Heatmaps, where each pixel represents the deviation between the smoothed fold changes observed experimentally (±*TEL1* ratio) for each combination of simulated interference window width (vertical axis) and variable trans interference strength (0–1, horizontal axis) across the sixteen *S. cerevisiae* chromosomes. Dark red and yellow pixels indicate areas of low (<0.2) and high (>0.5) deviation between the experimental and the simulations, respectively. Either 150 (left panel), 175 (middle panel) or 200 (right panel) DSBs were simulated per cell, with an independent number of cells iterated until a total of 10 million DSBs had been simulated.

**Supplementary Figure 5.**
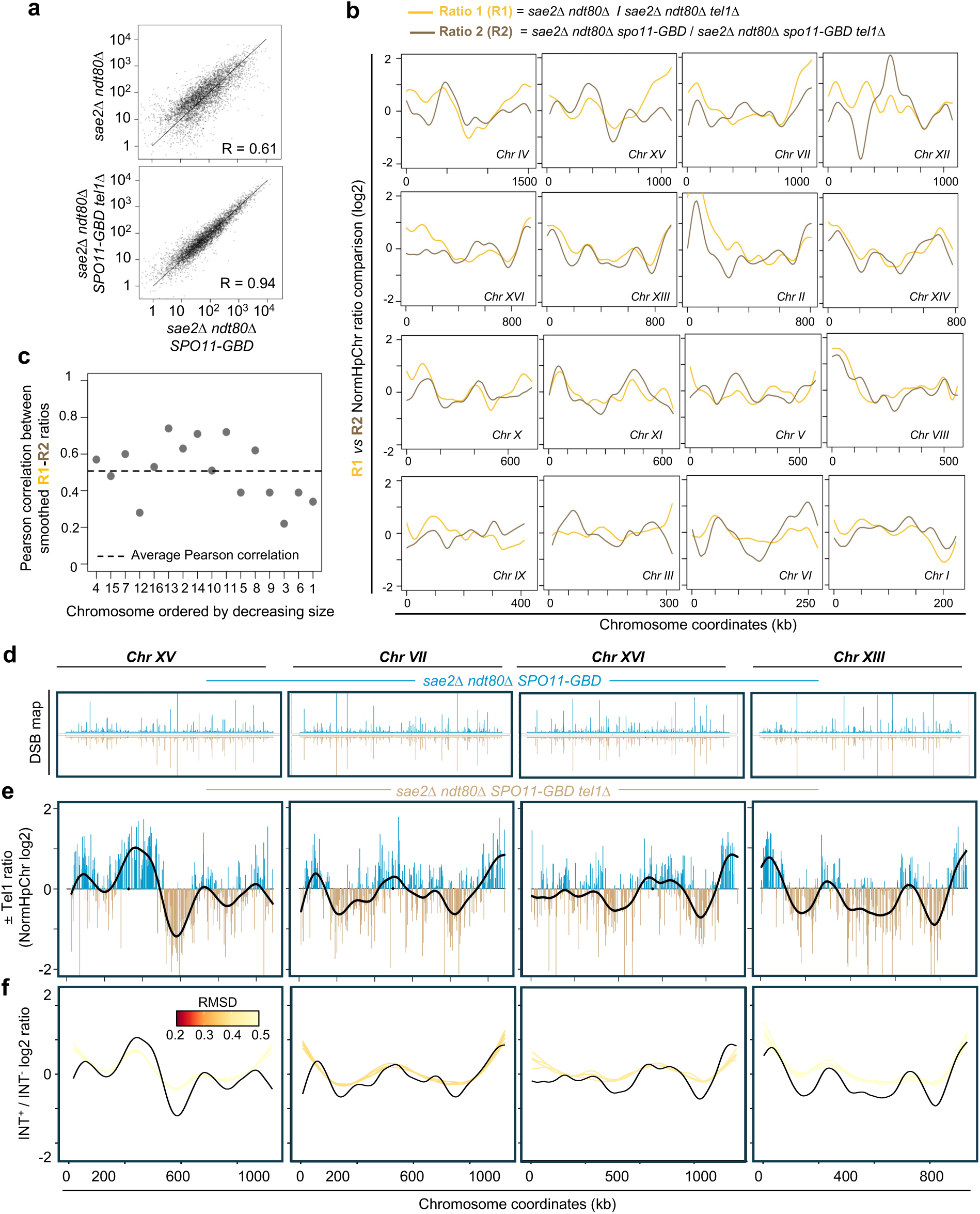
The impact of *SPO11-GBD* on Spo11-DSB distributions. **a**, Pearson correlation in hotspot strength between the indicated strains. **b**, Comparison between the log_2_ ratio of relative Spo11 hotspot intensities (NormHpChr) ±*TEL1* in the *SPO11* wild type (Ratio 1) and *SPO11-GBD* (Ratio 2) backgrounds across the sixteen *S. cerevisiae* chromosomes. **c**, Pearson correlation between *SPO11* wild type (R1) and *SPO11-GBD* (R2) smoothed log_2_ ratio of relative Spo11 hotspot intensities ±*TEL1* across the sixteen *S. cerevisiae* chromosomes. The dotted line represents the average Pearson correlation across all chromosomes. **d**, Visualization of the relative Spo11 hotspot intensities (NormHpChr) on the indicated chromosomes in the indicated strains. **e**, Ratio of relative Spo11 hotspot intensities ±*TEL1* on the indicated chromosomes in the *SPO11-GBD* system. Values above zero indicate a higher DSB frequency in the presence of Tel1 and below zero a higher DSB frequency in the absence of Tel1. Fold change was smoothed to highlight the spatial trend effect of *TEL1* deletion (black line) in the *SPO11-GBD* system. **f,** Representation of the simulations (INT+/INT- log2 ratio) that best recapture the experimental *±TEL1* smoothed pattern (NormHpChr log_2_ ratio; curves coloured by RMSD) in the SPO11-GBD backgrounds on the indicated chromosomes.

**Supplementary Figure 6.**
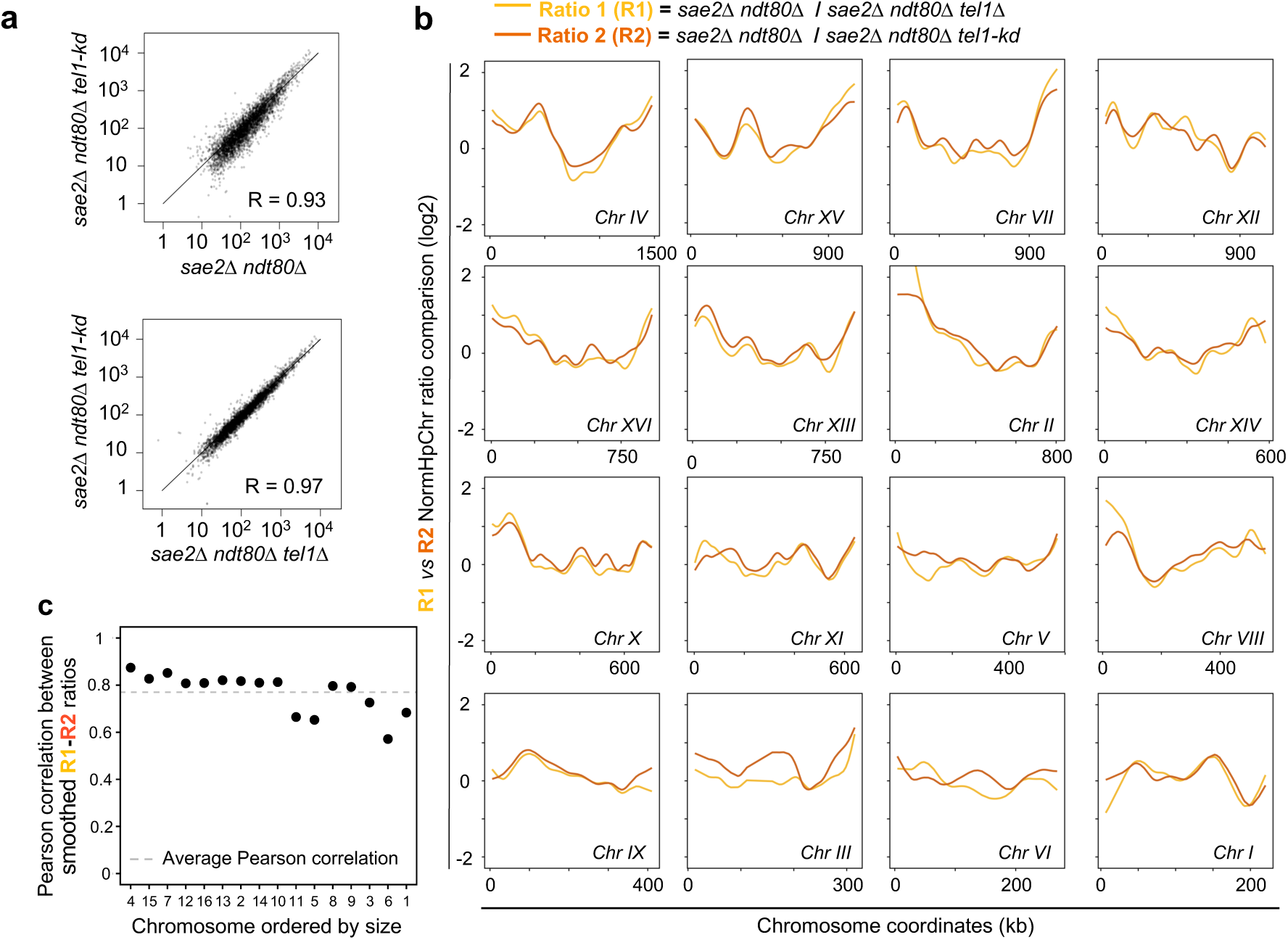
Comparison between effect of Tel1 and *tel1-kd* on DSB distributions. **a**, Pearson correlation in hotspot strength between the indicated strains. **b**, Comparison between the log_2_ ratio of relative Spo11 hotspot intensities (NormHpChr) ±*TEL1* (Ratio 1, R1) and *tel1-kd* (Ratio 2, R2) across the sixteen *S. cerevisiae* chromosomes. **c**, Pearson correlation between ±*TEL1* (R1) and *tel1-kd* (R2) smoothed log_2_ ratio of relative Spo11 hotspot intensities ±*TEL1* across the sixteen *S. cerevisiae* chromosomes. The dotted line represents the mean correlation value across all chromosomes.

**Supplementary Figure 7.**
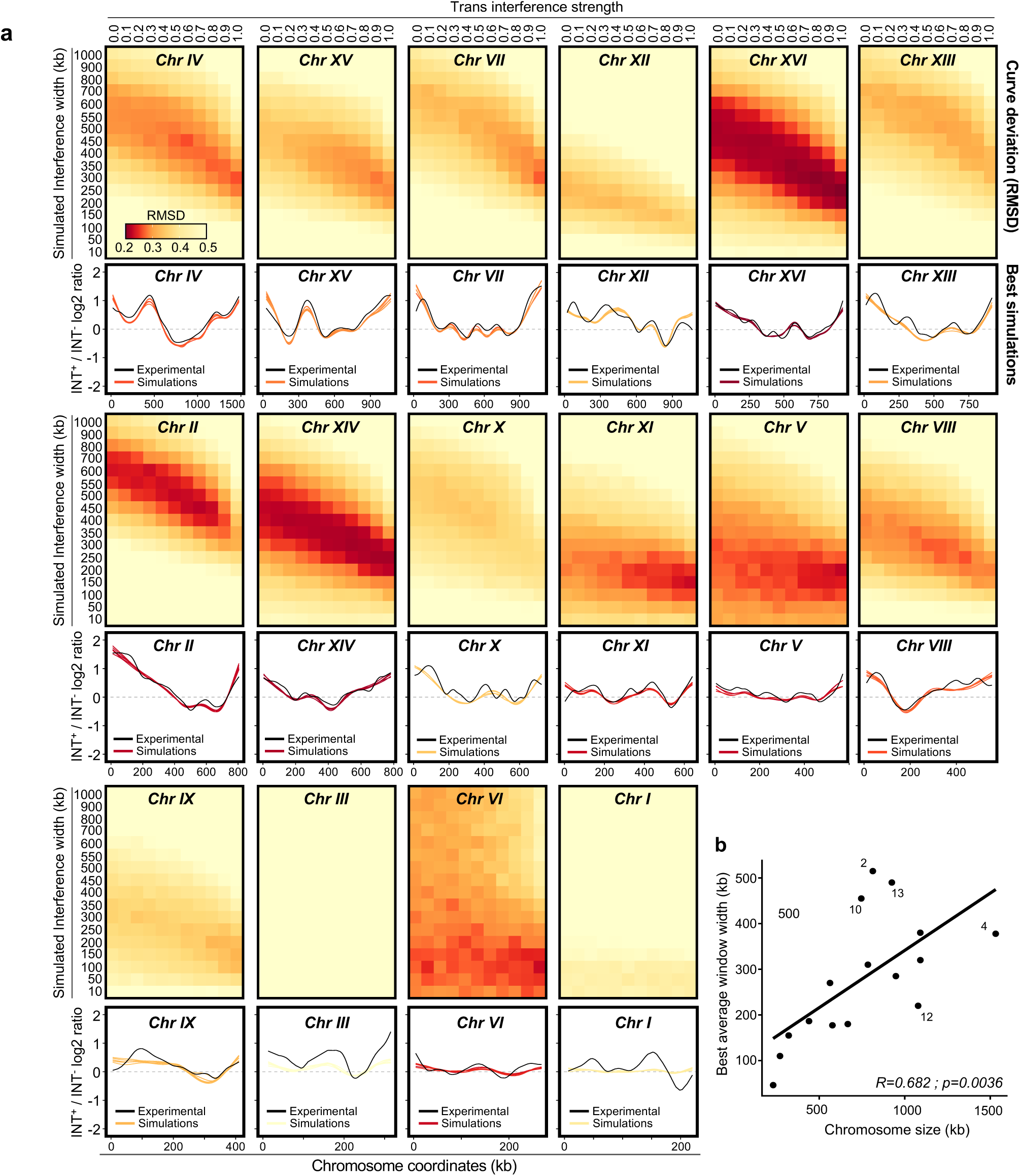
Simulation of DSB interference accurately recaptures the effect of *tel1-kd*. **a**, (Upper panels) Heatmaps, where each pixel represents the deviation between the smoothed fold changes observed experimentally (±*TEL1* ratio) for each combination of simulated interference window width (vertical axis) and variable trans interference strength (0–1, horizontal axis) across the sixteen *S. cerevisiae* chromosomes. Dark red and yellow pixels indicate areas of low (<0.2) and high (>0.5) deviation between the experimental and the simulations, respectively. (Lower panels) Comparison of the simulations with lowest RMSD (coloured by RMSD as in the upper panels) against the ±*tel1kd* experimental smoothed pattern (black line). 175 DSBs were simulated per cell. **b**, Correlation between the window width of best fitting simulations (mean of the 10 simulations with the lowest RMSD) against chromosome size.

**Supplementary Figure 8.**
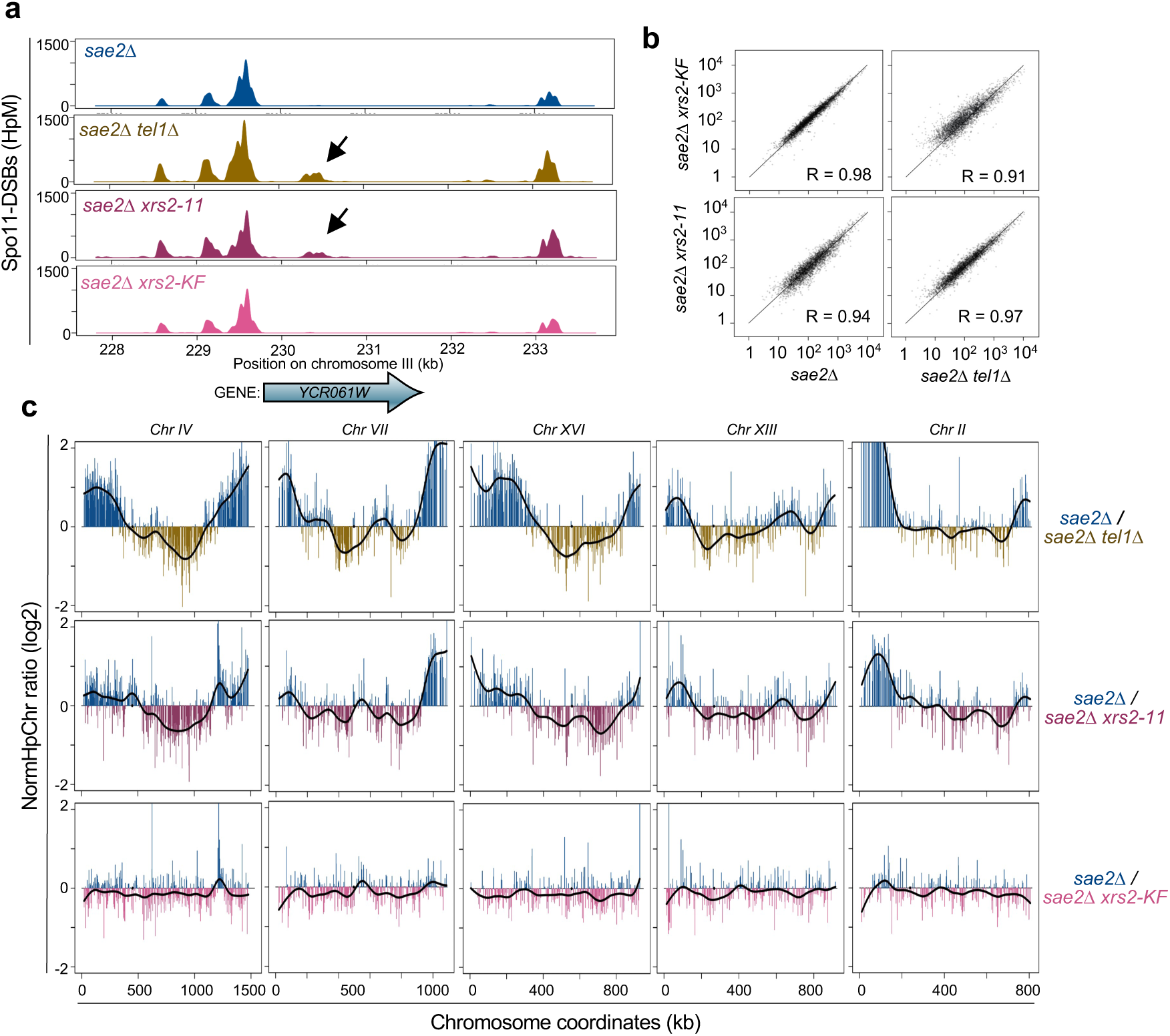
The Xrs2 C-terminal domain is required to mediate DSB interference. **a**, Visualization of the relative Spo11 hotspot intensities (HpM) in the *YCR061W* hotspot on chromosome III in the indicated strains. The black arrows indicate an adjacent hotspot that arises in the absence of Tel1. **b**, Pearson correlation of hotspot strength between the indicated strains. **c**, Ratio of relative Spo11 hotspot intensities (NormHpChr log2 ratio) ±*TEL1* (top panels), ±*xrs2-11* (middle panels) or ±*xrs2-KF* (bottom panels) for the indicated chromosomes. Values above zero indicate a higher DSB frequency in the presence of Tel1 (top panels) or Xrs2 wild type (middle and bottom panels) and below zero a higher DSB frequency in the absence of Tel1 (top panels) or in the presence of *xrs2-11* (middle panels) or *xrs2-KF* (bottom panels) constructs. Fold change was smoothed to highlight the spatial trend effect of *TEL1* deletion or the *xrs2* mutants (black line).

**Supplementary Figure 9.**
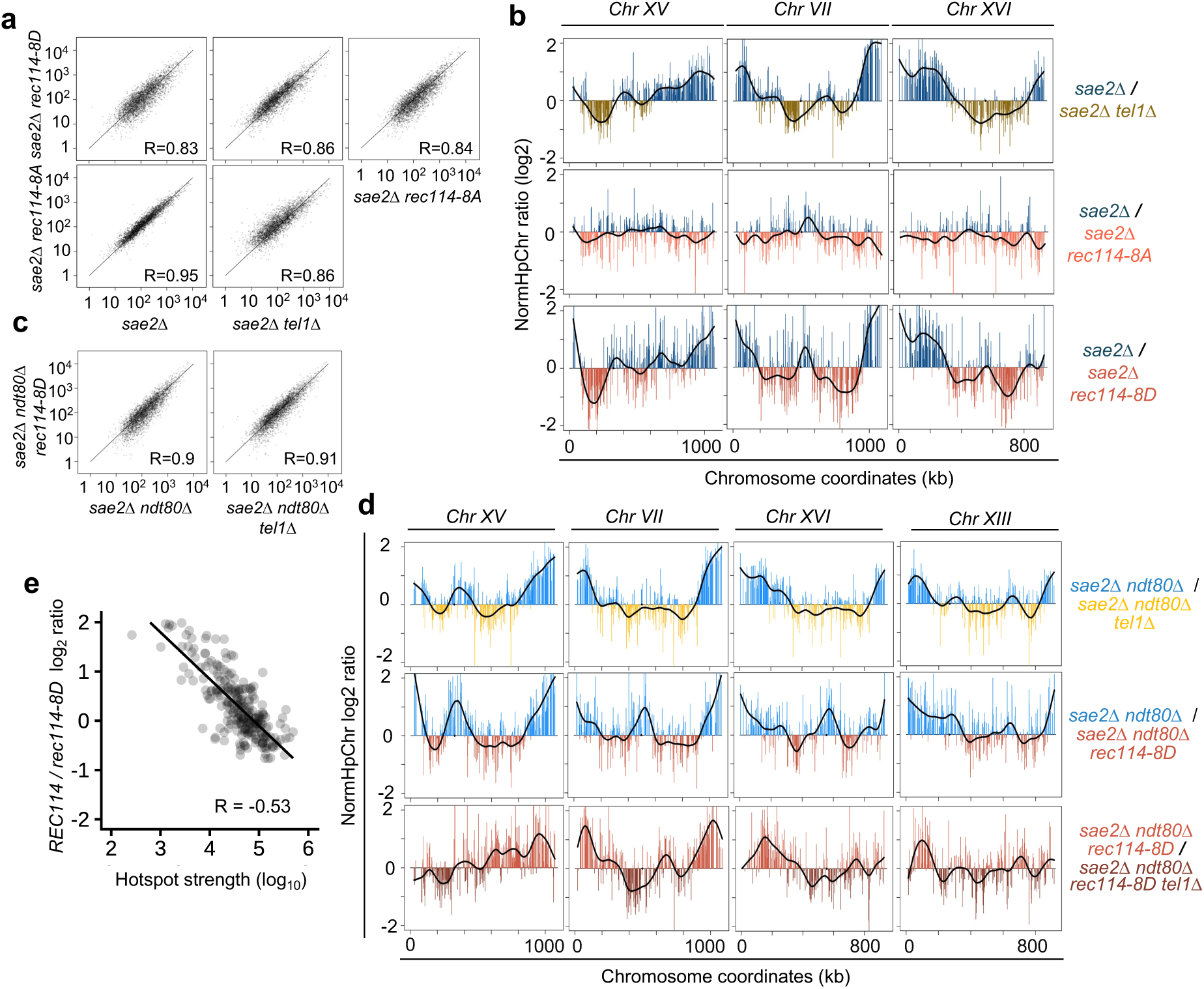
Effects of Rec114 mutants on DSB distributions across *S. cerevisiae* chromosomes. **a**, Pearson correlation in hotspot strength between the indicated strains. **b**, Ratio of relative Spo11 hotspot intensities (NormHpChr log2 ratio) ±*TEL1* (top panels), ±*rec114-8A* (middle panels) or ±*rec114- 8D* (bottom panels) on the indicated chromosomes. Values above zero indicate a higher DSB frequency in the presence of Tel1 (top panels) or Rec114 wild type (middle and bottom panels) and below zero a higher DSB frequency in the absence of Tel1 (top panels) or in the presence of *rec114-8A* (middle panels) or *rec114-8D* (bottom panels) constructs in the *sae2*Δ background. Fold change was smoothed to highlight the spatial trend effect of *TEL1* deletion or the *rec114-8A/8D* mutants (black line). **c**, Pearson correlation in hotspot strength between the indicated strains. **d**, Ratio of relative Spo11 hotspot intensities (NormHpChr log_2_ ratio) ±*TEL1* (top and bottom panels) or ±*rec114-8D* (middle panels) on the indicated chromosomes in the *sae2*Δ *ndt80*Δ background. Values above zero indicate a higher DSB frequency in the presence of Tel1 (top and bottom panels) or Rec114 wild type (middle panels) and below zero a higher DSB frequency in the absence of Tel1 (top and bottom panels) or in the presence of *rec114-8D* (middle panels) constructs. Fold change was smoothed to highlight the spatial trend effect of *TEL1* deletion or the *rec114-8D* mutant (black line). **e**, Pearson correlation between hotspot intensities across all chromosomes and *REC114*/*rec114-8D* log_2_ ratio.

**Supplementary Table 1.**
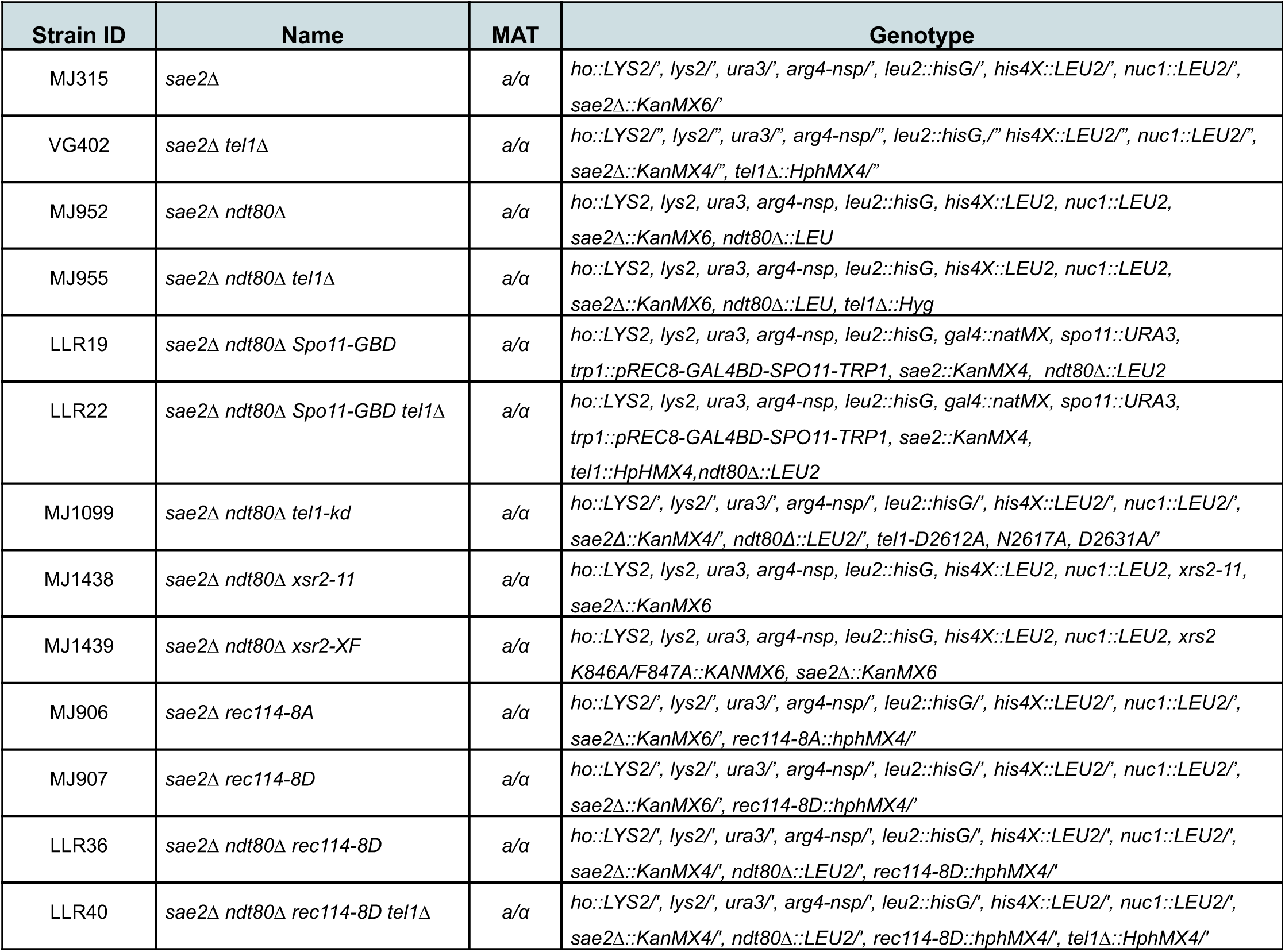
*S. cerevisiae* strains used in this study. All strains are isogenic from the SK1 background.

| Strain ID | Name | MAT | Genotype |
| --- | --- | --- | --- |
| MJ315 | *sae2*Δ | a/α | *ho::LYS2*′, *lys2*′, *ura3*′, *arg4-nsp*′, *leu2::hisG*′, *his4X::LEU2*′, *nuc1::LEU2*′, *sae2*Δ::KanMX6′ |
| VG402 | *sae2*Δ *tel1*Δ | a/α | *ho::LYS2*′, *lys2*′, *ura3*′, *arg4-nsp*′, *leu2::hisG*′, *his4X::LEU2*′, *nuc1::LEU2*′, *sae2*Δ::KanMX4′, *tel1*Δ::HphMX4′ |
| MJ952 | *sae2*Δ *ndt80*Δ | a/α | *ho::LYS2*, *lys2*, *ura3*, *arg4-nsp*, *leu2::hisG*, *his4X::LEU2*, *nuc1::LEU2*, *sae2*Δ::KanMX6, *ndt80*Δ::LEU |
| MJ955 | *sae2*Δ *ndt80*Δ *tel1*Δ | a/α | *ho::LYS2*, *lys2*, *ura3*, *arg4-nsp*, *leu2::hisG*, *his4X::LEU2*, *nuc1::LEU2*, *sae2*Δ::KanMX6, *ndt80*Δ::LEU, *tel1*Δ::Hyg |
| LLR19 | *sae2*Δ *ndt80*Δ *Spo11*-GBD | a/α | *ho::LYS2*, *lys2*, *ura3*, *arg4-nsp*, *leu2::hisG*, *gal4::natMX*, *spo11::URA3*, *trp1::pREC8-GAL4BD-SPO11-TRP1*, *sae2::KanMX4*, *ndt80*Δ::LEU2 |
| LLR22 | *sae2*Δ *ndt80*Δ *Spo11*-GBD *tel1*Δ | a/α | *ho::LYS2*, *lys2*, *ura3*, *arg4-nsp*, *leu2::hisG*, *gal4::natMX*, *spo11::URA3*, *trp1::pREC8-GAL4BD-SPO11-TRP1*, *sae2::KanMX4*, *tel1::HpHMX4*, *ndt80*Δ::LEU2 |
| MJ1099 | *sae2*Δ *ndt80*Δ *tel1*-kd | a/α | *ho::LYS2*′, *lys2*′, *ura3*′, *arg4-nsp*′, *leu2::hisG*′, *his4X::LEU2*′, *nuc1::LEU2*′, *sae2*Δ::KanMX4′, *ndt80*Δ::LEU2′, *tel1*-D2612A, N2617A, D2631A′ |
| MJ1438 | *sae2*Δ *ndt80*Δ *xsr2*-11 | a/α | *ho::LYS2*, *lys2*, *ura3*, *arg4-nsp*, *leu2::hisG*, *his4X::LEU2*, *nuc1::LEU2*, *xrs2*-11, *sae2*Δ::KanMX6 |
| MJ1439 | *sae2*Δ *ndt80*Δ *xsr2*-XF | a/α | *ho::LYS2*, *lys2*, *ura3*, *arg4-nsp*, *leu2::hisG*, *his4X::LEU2*, *nuc1::LEU2*, *xrs2* K846A/F847A::KANMX6, *sae2*Δ::KanMX6 |
| MJ906 | *sae2*Δ *rec114*-8A | a/α | *ho::LYS2*′, *lys2*′, *ura3*′, *arg4-nsp*′, *leu2::hisG*′, *his4X::LEU2*′, *nuc1::LEU2*′, *sae2*Δ::KanMX6′, *rec114*-8A::hphMX4′ |
| MJ907 | *sae2*Δ *rec114*-8D | a/α | *ho::LYS2*′, *lys2*′, *ura3*′, *arg4-nsp*′, *leu2::hisG*′, *his4X::LEU2*′, *nuc1::LEU2*′, *sae2*Δ::KanMX6′, *rec114*-8D::hphMX4′ |
| LLR36 | *sae2*Δ *ndt80*Δ *rec114*-8D | a/α | *ho::LYS2*′, *lys2*′, *ura3*′, *arg4-nsp*′, *leu2::hisG*′, *his4X::LEU2*′, *nuc1::LEU2*′, *sae2*Δ::KanMX4′, *ndt80*Δ::LEU2′, *rec114*-8D::hphMX4′ |
| LLR40 | *sae2*Δ *ndt80*Δ *rec114*-8D *tel1*Δ | a/α | *ho::LYS2*′, *lys2*′, *ura3*′, *arg4-nsp*′, *leu2::hisG*′, *his4X::LEU2*′, *nuc1::LEU2*′, *sae2*Δ::KanMX4′, *ndt80*Δ::LEU2′, *rec114*-8D::hphMX4′, *tel1*Δ::HphMX4′ |

**Supplementary Table 2.**
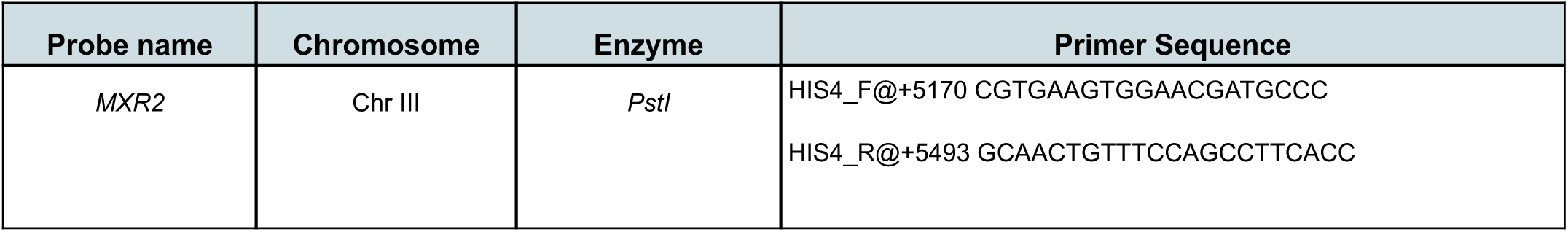
Oligonucleotides used in this study for Southern Blotting.

**Supplementary Table 3.**
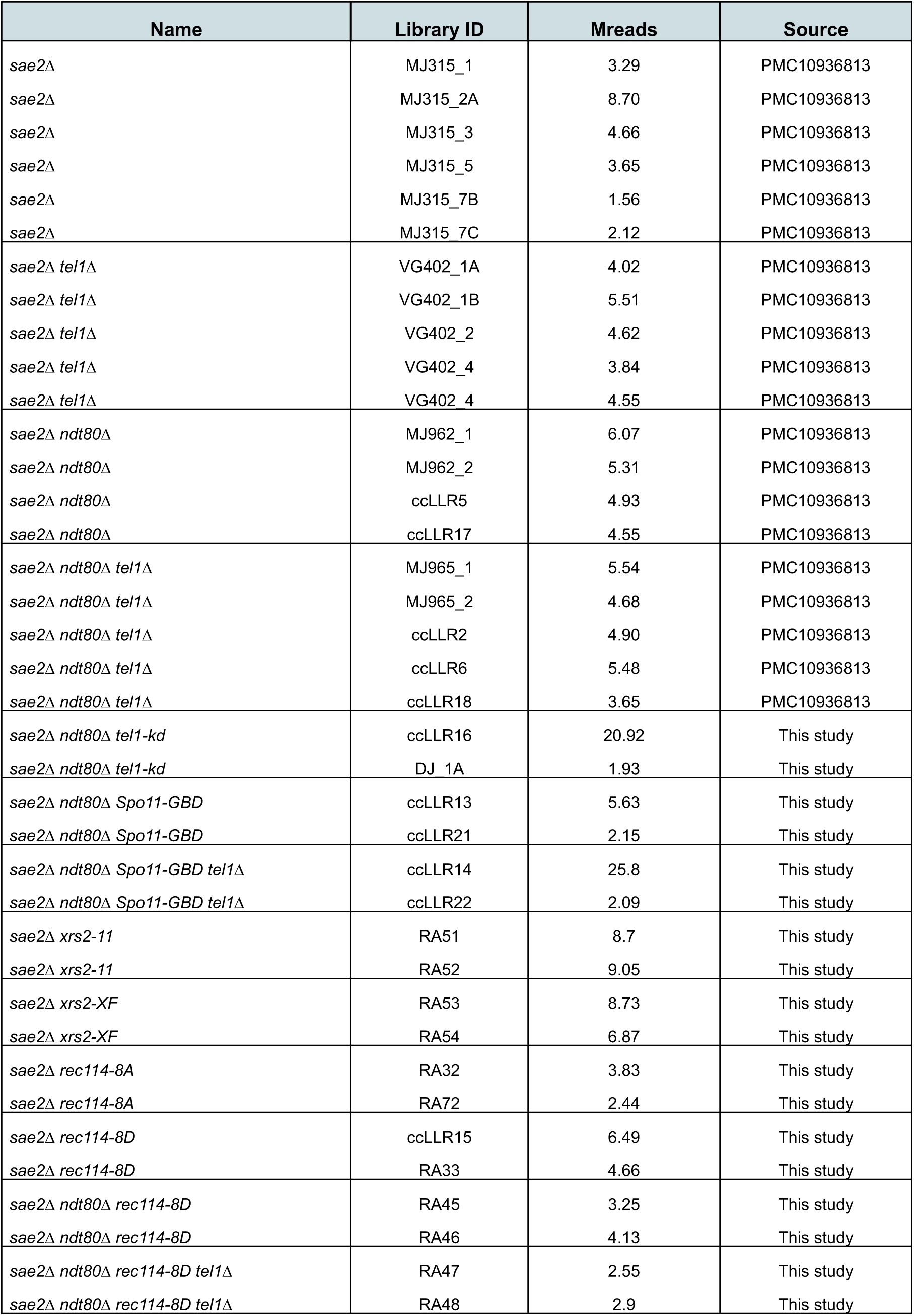
CC-seq libraries used in this study. Individual CC-seq libraries were prepared from biological sample replicates as indicated and sequenced using paired-end Illumina sequencing to the indicated read depth (Mreads, Million mapped reads). Libraries from the same genotype were averaged with equal weighting to generate the final averaged datasets used in all analyses.

| Name | Library ID | Mreads | Source |
| --- | --- | --- | --- |
| *sae2Δ* | MJ315_1 | 3.29 | PMC10936813 |
| *sae2Δ* | MJ315_2A | 8.70 | PMC10936813 |
| *sae2Δ* | MJ315_3 | 4.66 | PMC10936813 |
| *sae2Δ* | MJ315_5 | 3.65 | PMC10936813 |
| *sae2Δ* | MJ315_7B | 1.56 | PMC10936813 |
| *sae2Δ* | MJ315_7C | 2.12 | PMC10936813 |
| *sae2Δ tel1Δ* | VG402_1A | 4.02 | PMC10936813 |
| *sae2Δ tel1Δ* | VG402_1B | 5.51 | PMC10936813 |
| *sae2Δ tel1Δ* | VG402_2 | 4.62 | PMC10936813 |
| *sae2Δ tel1Δ* | VG402_4 | 3.84 | PMC10936813 |
| *sae2Δ tel1Δ* | VG402_4 | 4.55 | PMC10936813 |
| *sae2Δ ndt80Δ* | MJ962_1 | 6.07 | PMC10936813 |
| *sae2Δ ndt80Δ* | MJ962_2 | 5.31 | PMC10936813 |
| *sae2Δ ndt80Δ* | ccLLR5 | 4.93 | PMC10936813 |
| *sae2Δ ndt80Δ* | ccLLR17 | 4.55 | PMC10936813 |
| *sae2Δ ndt80Δ tel1Δ* | MJ965_1 | 5.54 | PMC10936813 |
| *sae2Δ ndt80Δ tel1Δ* | MJ965_2 | 4.68 | PMC10936813 |
| *sae2Δ ndt80Δ tel1Δ* | ccLLR2 | 4.90 | PMC10936813 |
| *sae2Δ ndt80Δ tel1Δ* | ccLLR6 | 5.48 | PMC10936813 |
| *sae2Δ ndt80Δ tel1Δ* | ccLLR18 | 3.65 | PMC10936813 |
| *sae2Δ ndt80Δ tel1-kd* | ccLLR16 | 20.92 | This study |
| *sae2Δ ndt80Δ tel1-kd* | DJ_1A | 1.93 | This study |
| *sae2Δ ndt80Δ Spo11-GBD* | ccLLR13 | 5.63 | This study |
| *sae2Δ ndt80Δ Spo11-GBD* | ccLLR21 | 2.15 | This study |
| *sae2Δ ndt80Δ Spo11-GBD tel1Δ* | ccLLR14 | 25.8 | This study |
| *sae2Δ ndt80Δ Spo11-GBD tel1Δ* | ccLLR22 | 2.09 | This study |
| *sae2Δ xrs2-11* | RA51 | 8.7 | This study |
| *sae2Δ xrs2-11* | RA52 | 9.05 | This study |
| *sae2Δ xrs2-XF* | RA53 | 8.73 | This study |
| *sae2Δ xrs2-XF* | RA54 | 6.87 | This study |
| *sae2Δ rec114-8A* | RA32 | 3.83 | This study |
| *sae2Δ rec114-8A* | RA72 | 2.44 | This study |
| *sae2Δ rec114-8D* | ccLLR15 | 6.49 | This study |
| *sae2Δ rec114-8D* | RA33 | 4.66 | This study |
| *sae2Δ ndt80Δ rec114-8D* | RA45 | 3.25 | This study |
| *sae2Δ ndt80Δ rec114-8D* | RA46 | 4.13 | This study |
| *sae2Δ ndt80Δ rec114-8D tel1Δ* | RA47 | 2.55 | This study |
| *sae2Δ ndt80Δ rec114-8D tel1Δ* | RA48 | 2.9 | This study |

